# Mechanical forces stimulate Golgi export

**DOI:** 10.1101/2025.09.02.673725

**Authors:** Chandini Bhaskar Naidu, Javier Vera Lillo, Eugenia Almacellas, Anabel-Lise Le Roux, Sabine Bardin, Nicolas Mateos, Jessica Angulo-Capel, Adam Wolowczyk, Yugo Terashima, Yuichi Wakana, Pere Roca-Cusachs, Maria F. Garcia-Parajo, Franck Perez, Bruno Goud, Jean-Baptiste Manneville, Stéphanie Miserey, Felix Campelo

**Affiliations:** Intracellular Transport: Engineering and Mechanisms Laboratory, Institut Curie, PSL Research University, Sorbonne Université, Centre National de la Recherche Scientifique, UMR 144, Paris, France; Laboratoire Matières et Systèmes Complexes, UMR 7057, CNRS and Université Paris Cité, CNRS, UMR7057, 10 rue Alice Domon et Léonie Duquet, F-75013, Paris cedex 13, France; ICFO-Institut de Ciencies Fotoniques, The Barcelona Institute of Science and Technology, Barcelona, Spain; Institute for Bioengineering of Catalonia (IBEC), The Barcelona Institute of Technology (BIST), Barcelona, Spain; School of Life Sciences, Tokyo University of Pharmacy and Life Sciences, Hachioji, Tokyo 192-0392, Japan; University of Barcelona, Barcelona, Spain; Institució Catalana de Recerca i Estudis Avançats (ICREA), Barcelona, Spain; Department of Medicine and Life Sciences (MELIS), Universitat Pompeu Fabra (UPF), Barcelona, Spain

**Author notes:** These authors contributed equally (listed in alphabetical order) (Chandini Bhaskar Naidu, Javier Vera Lillo). These authors contributed equally (Jean-Baptiste Manneville, Stéphanie Miserey, Felix Campelo).

## Abstract

Cells face diverse mechanical stimuli that vary with cell type, state, and pathological conditions. Mechanobiology investigates how cells sense and respond to these forces. While most work has focused on the cell surface and nucleus as primary mechanosensors, how intracellular organelles adapt to extracellular mechanical forces remains largely unknown. Here, we show that extracellular mechanical signals influence the secretory function of the Golgi apparatus. By subjecting adherent cells to mechanical challenges –cell spreading on different ligands, altered substrate stiffness, or equibiaxial strain–, we reveal that extracellular forces modulate Golgi-to-cell surface carrier biogenesis, thereby regulating exocytosis. Together with changes in Golgi membrane tension, we identify molecular determinants of the mechanotransduction pathway, including microtubule acetylation, diacylglycerol production, and protein kinase D activity. In turn, inhibition of Golgi export suppresses this mechanoresponse and causes impaired cell spreading. These findings uncover a bidirectional mechanotransduction axis in which extracellular mechanics tune Golgi secretory output, providing a framework for investigating organelle-based mechanoadaptation in physiology and disease.

## INTRODUCTION

Exocytosis enables cells to deliver molecules to their surface or secrete them into the extracellular space, thereby maintaining plasma membrane (PM) and extracellular matrix (ECM) homeostasis and organization and facilitating interactions with the environment. Newly synthesized secretory and membrane proteins are transported along intracellular routes from the endoplasmic reticulum (ER) through the Golgi complex and to the *trans*-Golgi network (TGN). At the TGN, cargos are sorted and packaged into transport carriers, which are rapidly directed to the PM along microtubule (MT) tracks. Upon carrier fusion with the cell surface, transmembrane proteins are incorporated into the PM, while soluble cargos are released into the extracellular space through exocytosis.

Similar to the highly heterogeneous organization of the PM, exocytic events are spatially regulated (Grigoriev et al., 2007, 2011; Yuan et al., 2015; Robinson et al., 1995; Huet-Calderwood et al., 2023; Fourriere et al., 2019; Stehbens et al., 2014), ensuring that secretion is precisely coordinated with cellular architecture and function. Previous studies have demonstrated that many exocytic events occur preferentially near focal adhesions (FAs) (Stehbens et al., 2014; Huet-Calderwood et al., 2017; Eisler et al., 2018; Lachuer et al., 2023; Huet-Calderwood et al., 2023; Fourriere et al., 2019). In particular, by combining the Retention Using Selective Hooks (RUSH) assay (Boncompain et al., 2012) –which synchronizes anterograde cargo transport– with a Selective Protein Immobilization (SPI) assay, we previously mapped the precise spatial organization of cargo arrival sites at the PM. Interestingly, our findings revealed that exocytosis does not occur randomly across the cell surface but rather at hotspots juxtaposed to FAs (Fourriere et al., 2019). Among the carriers involved in these targeted exocytic events are Golgi-derived, RAB6-positive carriers (Fourriere et al., 2019). RAB6 mediates multiple Golgi-to-PM export routes, including the trafficking of CARTS (carriers of the TGN to the cell surface), a class of RAB6- and protein kinase D (PKD)-dependent Golgi-to-PM transport carriers (Wakana et al., 2012).

FAs are dynamic, force-sensitive molecular platforms localized at the cell membrane. The core components of FAs are integrins, heterodimeric transmembrane receptors that connect the ECM with cytoplasmic components, enabling the transmission of extracellular signals into the cell. These outside-in signals control essential processes such as cell migration, tissue morphogenesis, and PM homeostasis (Kanchanawong and Calderwood, 2023). While substantial progress has been made in understanding how FAs (Kechagia et al., 2019) and the nucleus (Kirby and Lammerding, 2018; Cho et al., 2017) sense and transduce extracellular mechanical stimuli, it is far less understood whether and how intracellular organelles such as the Golgi reciprocally adapt their activity to these cues and thereby contribute to the overall cellular mechanoresponse. The aforementioned observation that membrane regions proximal to FAs act as hotspots for exocytosis suggests a spatial coordination of membrane trafficking and adhesion sites. Exocytosis delivers integrins, ECM components (such as collagens, fibronectin and fibrillin), and signaling molecules to the PM (Moreno-Layseca et al., 2019), thereby sustaining adhesion dynamics. While MTs have long been recognized to target FAs (Schmidt and Stehbens, 2024; Krylyshkina et al., 2003; Seetharaman and Etienne-Manneville, 2019; Kaverina et al., 1998), emerging evidence indicates that exocytosis near FAs is spatially orchestrated by MT guidance, RAB GTPase-dependent Golgi-to-PM trafficking, and, potentially, mechanochemical feedback loops. This spatial regulation may provide a mechanism for localizing membrane expansion, modulating membrane tension, and remodeling the ECM and PM proteome –processes critical for adhesion homeostasis during cell spreading and migration. Understanding these dynamics may also offer insights into how dysregulation of these processes contributes to pathological conditions such as cancer invasion.

While several mechanosensitive modules, such as FAs, fibrillar adhesions, and adherens junctions at the PM (Elosegui-Artola et al., 2018; De Pascalis and Etienne-Manneville, 2017), and the nuclear envelope (Thorpe and Lee, 2017; Amar et al., 2021; Jahed and Mofrad, 2019) have been well characterized, recent studies indicate that other organelles, such as the ER and lysosomes, can respond to mechanical forces (Phuyal and Baschieri, 2020; Rawal et al., 2025; Merta et al., 2025; Aceiton, 2025; Ren et al., 2025; Naughton et al., 2025). Moreover, mechanical forces applied to the PM can trigger Ca^2+^ release from the ER via an actomyosin-dependent mechanism (Kim et al., 2015). In the context of intracellular trafficking, neurite mechanical tension enhances active vesicular transport in neurons (Ahmed and Saif, 2014), while COPII-mediated transport from the ER to the Golgi apparatus is regulated by mechanical strain through interactions between the small G proteins Rac1 and Sar1 (Phuyal et al., 2022), and is sensitive to ER membrane viscosity (Jimenez-Rojo, 2025). The Golgi apparatus itself is mechanoresponsive: external forces generated by adhesion to the ECM influence lipid metabolism and Golgi mechanics (Romani et al., 2019), and cell–matrix adhesion controls Golgi organization and secretory output via an ARF1- and AXL-dependent pathway (Singh et al., 2018; Saha et al., 2026; Chakraborty et al., 2026; Joshi et al., 2026). Because the actomyosin cytoskeleton (Zilberman et al., 2011; Miserey-Lenkei et al., 2010) and MT networks (Fourriere et al., 2020) are closely associated with Golgi membranes, forces transmitted through the cytoskeleton can directly alter Golgi dynamics (Miserey-Lenkei et al., 2017). In addition, using optical-tweezers based microrheology, we showed that directly applying force to the Golgi apparatus delays actin-mediated membrane fission and thereby perturbs post-Golgi trafficking, indicating that Golgi membranes are mechanosensitive (Guet et al., 2014).

In this study, we address the impact of extracellular mechanical forces on post-Golgi carrier biogenesis. Our results show that conditions associated with impaired FA formation are accompanied by reduced Golgi-derived carrier biogenesis, whereas both substrate stiffening and mechanical stretching enhance Golgi export capabilities. Using Halo-Flipper probes, we further show that extracellular mechanical forces are accompanied by changes in Golgi membrane mechanical properties. This is paralleled with an increase in MT acetylation, changes in Golgi diacylglycerol (DAG) content, and PKD activity. In turn, inhibition of Golgi export suppresses this mechanoresponse, resulting in impaired cell spreading. Collectively, our data identifies the Golgi apparatus as a mechanoresponsive organelle, revealing that its export activity is modulated by extracellular mechanical stimuli and is crucial for efficient cell spreading and mechanoadaptation.

## RESULTS

### Golgi-derived carrier biogenesis scales with cell adhesion and FA formation

In non-polarized 2D cell cultures, we and others have shown that exocytic sites are not randomly distributed across the PM but preferentially localize close to FAs in a RAB6-dependent manner (Fourriere et al., 2019). RAB6 is a general regulator of post-Golgi carrier biogenesis, including CARTS (Wakana et al., 2012). However, whether CARTS exocytosis itself is spatially regulated with respect to FAs has not been directly tested. To address this, we mapped the delivery of the CARTS-specific secretory cargo Pancreatic Adenocarcinoma Upregulated Factor (PAUF). HeLa cells transiently co-expressing paxillin-GFP (FA marker) and mKate2-FM4-PAUF (CARTS cargo) were imaged by live-cell total internal reflection fluorescence (TIRF) microscopy. After exiting the Golgi apparatus, PAUF-positive carriers moved quasi-directionally through the cytoplasm (***Video 1***) and docked preferentially near FA-rich regions of the PM (***Fig. S1A***). At these sites, CARTS often paused for several seconds before a sudden loss of fluorescence, consistent with fusion with the PM (***Fig. S1A***, panel *(i)*). By contrast, outside FA areas, CARTS were absent or observed only transiently, without apparent pausing, suggesting that these carriers were likely in transit toward FA sites for exocytosis (***Fig. S1A***, panel *(ii)*). To further support this spatial relationship, we examined whether CARTS distribution is affected by depletion of ELKS, a cortical RAB6 effector previously shown to promote docking and fusion of RAB6-positive secretory carriers at exocytic hotspots near FAs (Fourriere et al., 2019). ELKS knockdown by siRNA (***Fig. S1B)*** led to an accumulation of PAUF-positive CARTS in cytoplasmic regions proximal to FAs (marked by paxillin) (***Fig. S1C, D***). This is consistent with previous observations for general RAB6 carriers and supports the idea that CARTS are preferentially targeted to FA-proximal exocytic hotspots (Fourriere et al., 2019). Finally, using a surface protein immobilization (SPI) assay at 40 min after cargo release from the ER, secreted PAUF accumulated at the periphery of the cell in proximity to FAs (***Fig. S1E, F***), similar to previous findings combining SPI and RUSH of other secretory cargoes (Fourriere et al., 2019).

To test whether integrin-mediated adhesion modulates Golgi-to-PM trafficking, we devised two complementary assays: *(i)* a combined spreading-secretion assay, and *(ii)* modulation of FAs by substrate stiffness. First, for the combined spreading-secretion assay (***Fig. 1A***), HeLa cells in suspension were plated on glass coated with fibronectin (FN) or poly-L-lysine (PLL). As previously described (Fourriere et al., 2019), PLL-seeded cells failed to assemble FAs, contrary to FN, which promoted FA formation (***Fig. S2A***). Cells were co-transfected with GFP-mem (a general PM marker, see Methods) and mKate2-FM4-PAUF to measure by confocal imaging both cell spreading (cell adhesion area) and CARTS biogenesis (number of CARTS per cell at 30 min after cargo release from the ER) (***Fig. 1A***, see Methods). On FN, cells exhibited larger adhesion areas and produced significantly more CARTS than on PLL (***Fig. 1B–D***). We note that adhesion area is distinct from total cell surface area and cell volume, and that spreading can be accompanied by changes in cell height and volume (Guo et al., 2017; Venkova et al., 2022). CARTS numbers in cells seeded on FN increased over time, peaking at 4 h post-seeding, whereas PLL-seeded cells showed persistently low CARTS counts even after 24 h (***Fig. 1C***). Likewise, the adhesion area of cells seeded on FN steadily expanded over 4 h, while it remained minimal and unaltered over 24 h in cells seeded on PLL (***Fig. 1D***). Similar results were obtained when cells were gently lifted with EDTA prior to the spreading-secretion assay, as compared to trypsinization, indicating that the observed differences were not attributable to trypsin-mediated cleavage of cell-surface proteins or trypsin-induced signaling (***Fig. S2B***). Importantly, CARTS number scaled linearly with adhesion area under all conditions, resulting in a constant number of CARTS per adhesion area (***Fig. 1E, F***). These results suggest that adhesion-dependent cell states characterized by increased adhesion area associate with enhanced Golgi export capacity in a commensurate manner.

**Figure 1.**
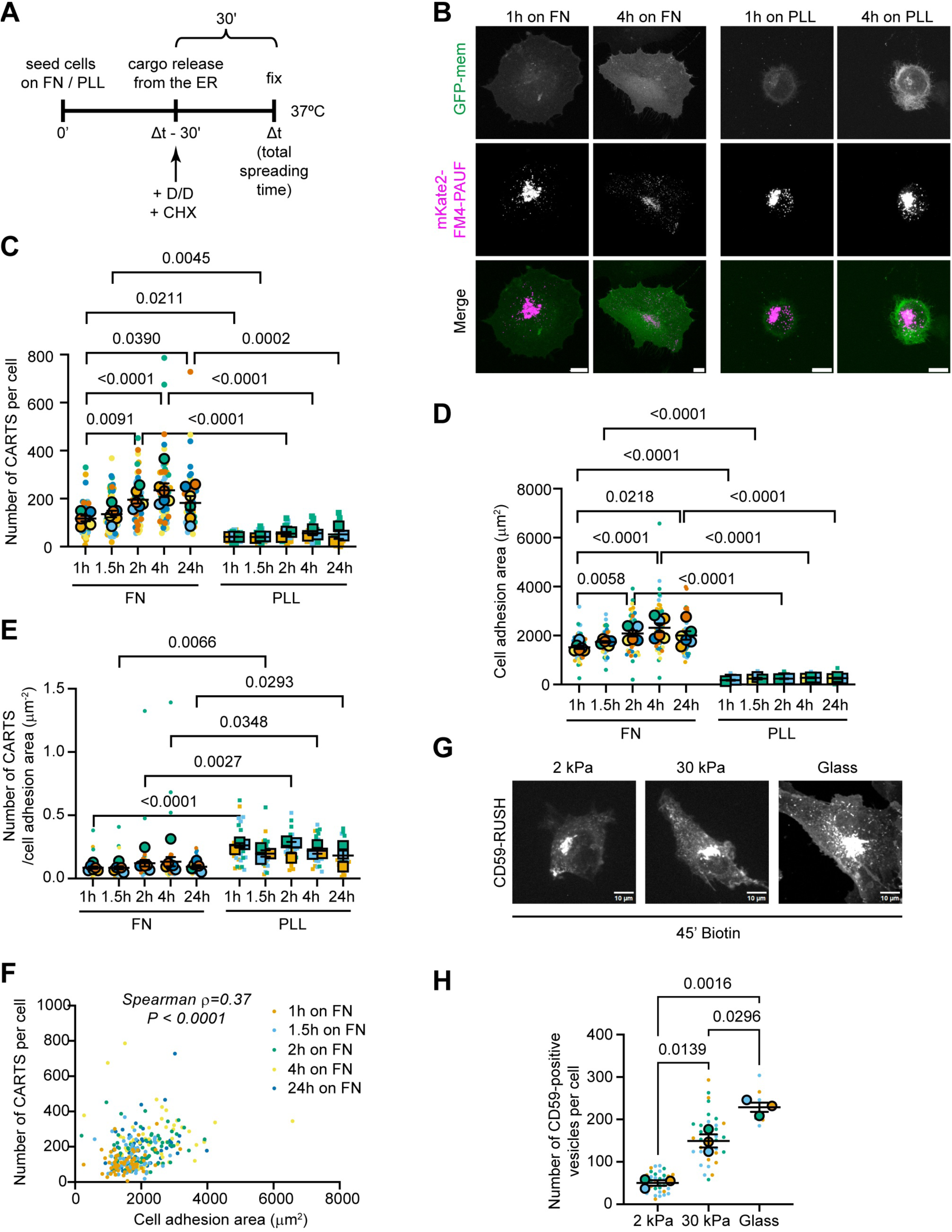
Golgi-derived carrier biogenesis scales with cell adhesion and FA formation. (**A**) Schematic representation of the spreading assay. D/D is D/D solubilizer, which dissolves mKate2-FM4-PAUF aggregates, allowing synchronized cargo release from the ER; CHX is cycloheximide. (**B**) Representative images of fixed HeLa cells transfected with GFP-mem and the mKate2-FM4-PAUF, seeded over FN or PLL, and acquired by confocal microscopy after the indicated times. Scale bars, 10 µm. (**C-E**) SuperPlots showing individual cell measurements (small, light-colored symbols; N∼10 per biological replicate) and the mean value for each independent biological replicate (larger, black-outlined circles; n=6 on FN, n=3 on PLL). Each color represents a different experimental replicate. Plots represent (**C**) the number of CARTS per cell, (**D**) the cell adhesion area, and (**E**) the number of CARTS per cell adhesion area. Repeated-measures 2-way ANOVA tests were performed, and P values were obtained using Tukey’s post-hoc multiple comparison test. (**F**) Correlation plot showing the number of CARTS per cell and the corresponding cell adhesion area per each measured individual cell across all tested conditions (see legend, cells seeded on FN only). Normality of both variables was tested using the Shapiro-Wilk test, and the results showed non-normality. Each point represents an individual cell. A nonparametric Spearman correlation analysis was performed, with the correlation coefficient being ρ=0.37 and P value < 0.0001. (**G**) Spinning disk confocal microscopy images of fixed HeLa cells stably expressing RUSH-EGFP-CD59 plated on FN-coated glass coverslips, or on stiff (30 kPa) or soft (2 kPa) FN-coated PAA gels. 4h after plating them, cells were incubated for 45 minutes with biotin before fixation to specifically visualize CD59-positive post-Golgi carriers. (**H**) SuperPlot showing quantification of the number of CD59-positive vesicles per cell in cells seeded on substrates of various stiffness as described in (**G**), and a repeated-measures 1-way ANOVA test was performed using Tukey’s post-hoc multiple comparison test (N=2–10, n=3).

Second, we performed a substrate stiffness assay to further manipulate FA assembly, as the number of FAs decreases with decreasing substrate stiffness, resulting in minimal FA formation on soft substrates (Pelham and Wang, 1997; Barber-Perez et al., 2020). HeLa cells were plated for 4 h on FN-coated polyacrylamide (PAA) gels of defined stiffness (2 kPa for soft gels and 30 kPa for stiff gels) or on glass, and FAs were visualized by immunofluorescence microscopy using vinculin staining (***Fig. S2C***). Cells on soft substrates spread poorly and formed almost no vinculin-positive FAs, whereas cells on stiff substrates displayed robust spreading and abundant FAs (***Fig. S2A,*** left panels and ***S2C***). We then assessed Golgi export using the RUSH assay, which enables synchronized trafficking of selected cargos (Boncompain et al., 2012). HeLa cells stably expressing RUSH-EGFP-CD59 (a RAB6-dependent GPI-anchored protein) were plated on soft (2 kPa PAA gels) and stiff (30 kPa PAA gels or glass) FN-coated substrates for 4 hours. Post-Golgi carriers were quantified at 45 min after biotin addition (***Fig. 1G***). Cells on stiff substrates exhibited ∼150–200 carriers per cell, compared with ∼50 carriers per cell on soft substrates (***Fig. 1H***). To directly assess whether active β_1_integrin levels at the basal membrane scale with this response, we stained cells using the active conformation-specific 9EG7 antibody (Bazzoni et al., 1995). While active β_1_ integrin levels at the basal cell surface tended to be higher on FN-coated glass than on 2 kPa or 30 kPa substrates, this difference was not statistically significant (***Fig. S2D, E***). By contrast, FA area was strongly reduced on 2 kPa or 30 kPa substrates as compared to glass, consistent with reduced spreading and lower carrier abundance under these conditions (***Fig. S2F***). This is in line with previous work showing that substrate stiffness regulates adhesion organization, including the elongation of active α_5_β_1_-integrin-positive fibrillar adhesions (Barber-Perez et al., 2020). Overall, our data support the hypothesis that adhesion-dependent cell states characterized by robust spreading and FA formation trigger an adaptive response in the secretory pathway.

### Mechanical stretch enhances Golgi export capabilities

Cell adhesion and spreading on integrin-activating substrates generate mechanical forces via integrin engagement and actomyosin contractility (Geiger et al., 2009). We have found that this scales with increased Golgi export capacity (***Fig. 1***). To test whether a distinct external mechanical cue produces similar effects, we applied sustained isotropic mechanical strain on cells using an equibiaxial stretching device (***Fig. 2A***) (see Methods; (Le Roux et al., 2025)), a mechanical stimulus that mimics ECM stretching. HeLa cells expressing PAUF-mRFP were seeded on FN-coated stretchable PDMS membranes and allowed to adhere for 1 h at 37°C. To synchronize cargo at the TGN, cells were then shifted to 20°C for 2h in the presence of cycloheximide (CHX). After shifting back to 37°C to resume Golgi export, cells were either left unstretched or subjected to a 15% isotropic strain for 30 min before fixation (***Fig. 2B***). Widefield imaging of the PAUF-mRFP signal revealed that mechanical stretch increased the number of cytoplasmic CARTS by ∼40% compared with unstretched controls (***Fig. 2C, D***). Measurements of cell adhesion area showed a variable increase (∼12% on average) upon stretch, which did not reach statistical significance (***Fig. 2E***). This may reflect dynamic remodeling and partial relaxation of the adhesion area during the 30 min sustained stretch. In addition, the number of CARTS per adhesion area showed a slight but not statistically significant increase (***Fig. 2F***). These data suggest that although cells rapidly remodel their adhesion footprint under sustained stretch, Golgi-derived carriers remain elevated at the end of the 30 min sustained stretch period. Stretch-induced changes in FA tension and/or mechanotransduction signaling may also contribute to the observed phenotype.

**Figure 2.**
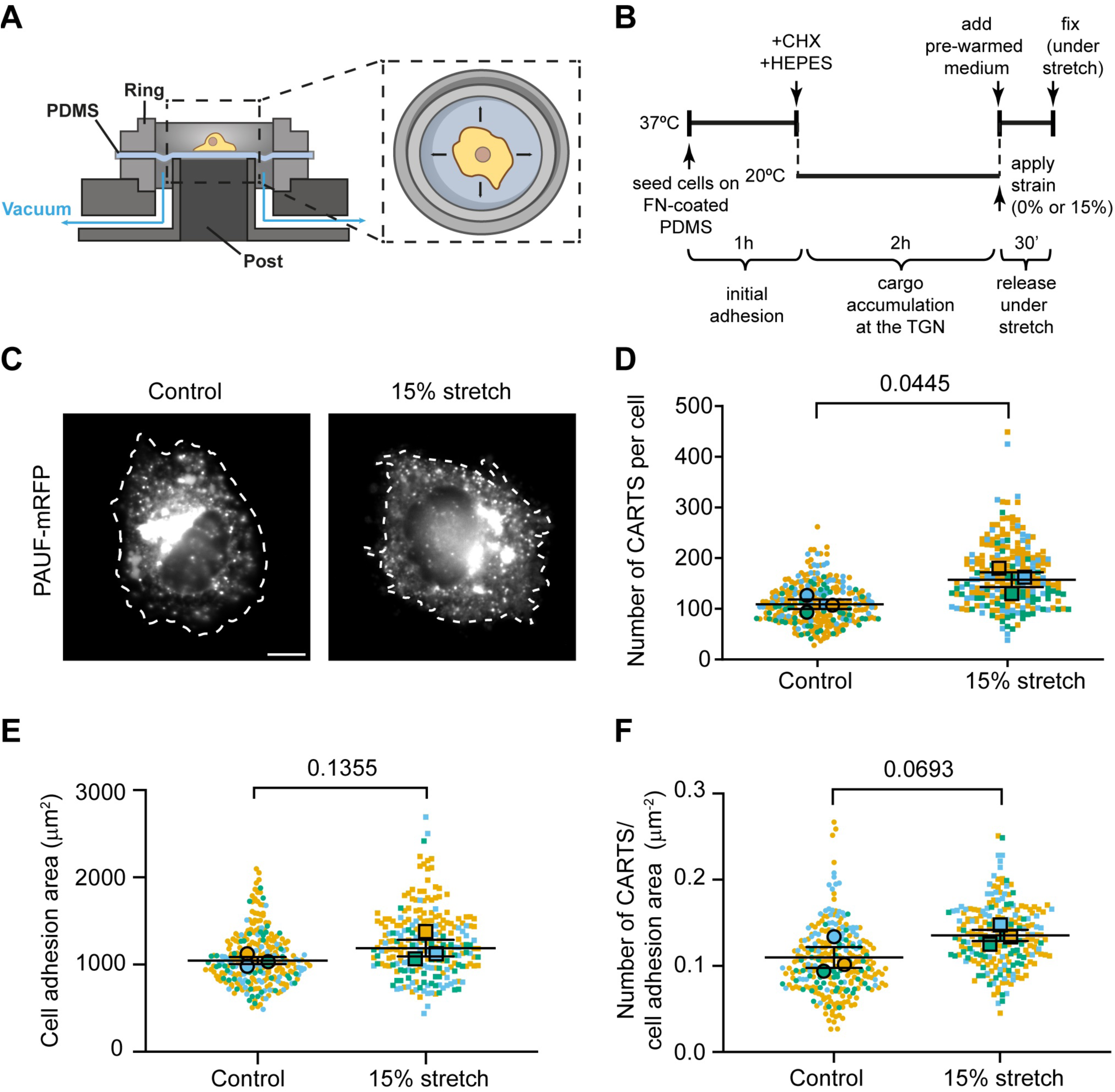
Mechanical strain application on cells increases their secretory capabilities. (**A**) Schematics of the stretch system used to apply a mechanical strain on cells. (**B**) Schematic representation of the pipeline followed in the stretching experiments. (**C**) Representative widefield images of fixed HeLa cells transfected with PAUF-RFP, seeded over FN-coated PDMS membranes, and subjected to no external forces or 15% equibiaxial stretch for 30 min. Cell contours are indicated by dashed white curves. Scale bar, 10 µm. (**D-F**) SuperPlots showing the indicated cell measurements (small, light-colored symbols; N>45 per biological replicate) and the mean value for each independent biological replicate (larger, black-outlined circles; n=3). Each color represents a different experimental replicate. Two-sided parametric ratio paired t-test was used. The P values are indicated in the plot.

### Mechanical forces impact Golgi membrane tension

Our previous work using optical tweezers demonstrated that forces applied directly on Golgi membranes alter their mechanical properties, and that actin depolymerization decreases Golgi rigidity (Guet et al., 2014). More recently, fluorescence lifetime imaging microscopy (FLIM)-based sensors of the Flipper family, such as Halo-Flipper, have been developed as lipid packing and membrane tension reporters (Strakova et al., 2019; Colom et al., 2018). Flippers respond to lipid packing, and thereby membrane tension, by shifting between planar and twisted conformations, which can be quantified by FLIM. We therefore tested whether extracellular mechanical forces alter Golgi tension using Halo-Flipper. First, to validate the use of Halo-Flipper at Golgi membranes, we exploited the effect of Latrunculin A on actin depolymerization, which is expected to reduce Golgi tension. RPE1-ManII-Halo stably expressing cells treated with low doses of Latrunculin A displayed compaction of the Golgi apparatus, as previously shown (Egea et al., 2006), and a significant decrease in Halo-Flipper fluorescence lifetime at the Golgi (***Fig. 3A, B***), consistent with reduced membrane tension. These results align with our previous optical tweezers measurements (Guet et al., 2014) and confirm that Halo-Flipper can report changes in Golgi membrane mechanical properties.

**Figure 3:**
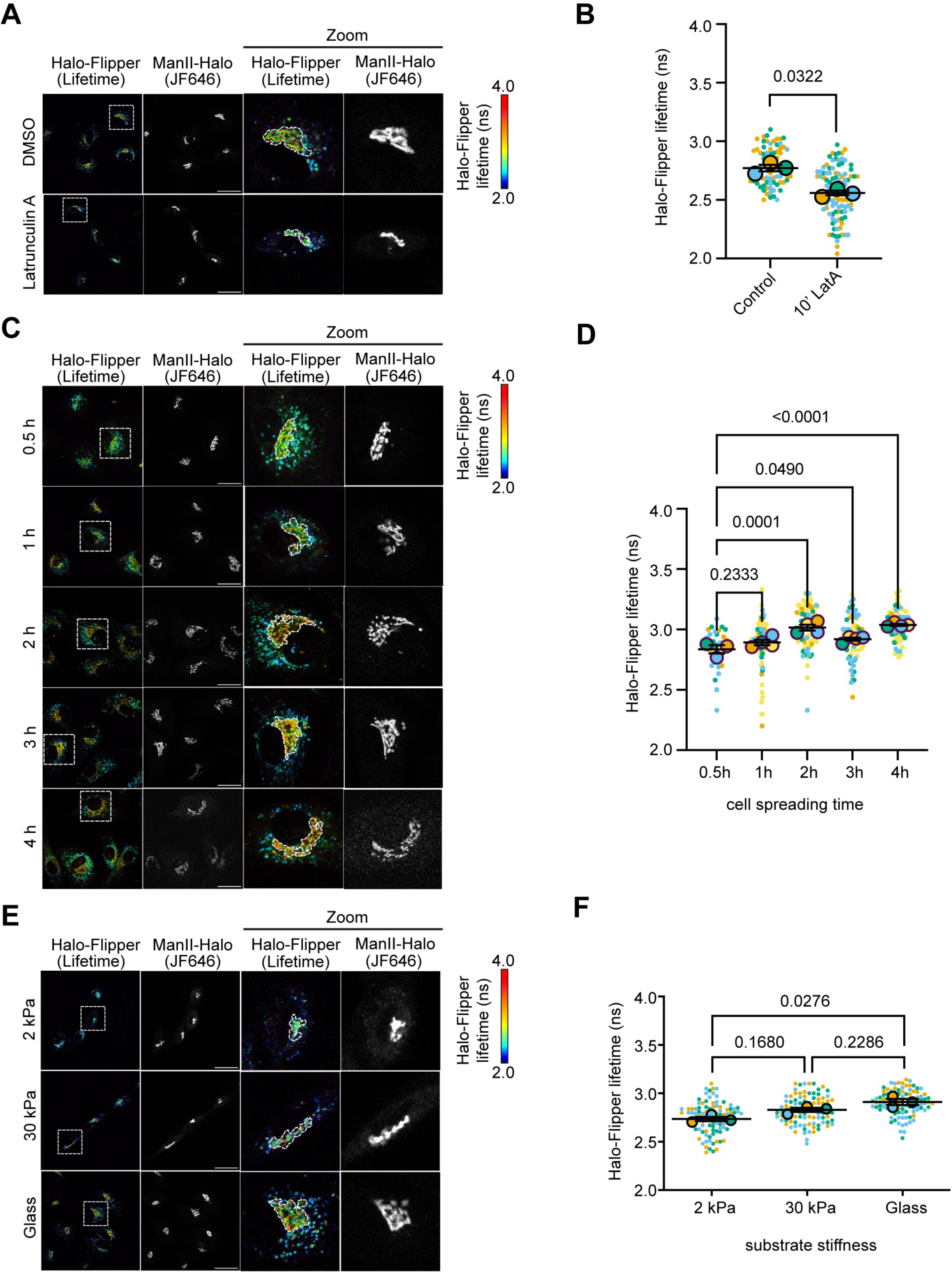
Golgi membrane tension responds to mechanical forces. (**A**) Images of live RPE1-ManII-Halo stably expressing cells before and after treatment with 50 nM Latrunculin A. The FLIM signal is displayed. The Golgi apparatus was post-labelled using JF-646. Higher magnification is shown on the right. (**B**) SuperPlot of FLIM value expressed as fluorescence lifetime in ns was quantified for each experimental condition (n=3 replicates; N>15 cells per replicate). A two-sided parametric ratio paired t-test was used. The P value is indicated in the plot. (**C**) Images of live RPE1-ManII-Halo stably expressing cells spread for 30 min to 4 h on FN-coated glass coverslips. The Flipper average lifetime obtained by FLIM is displayed (color scale). To specifically measure FLIM at the Golgi, ManII-Halo was post-labeled with JF-646 (intensity in gray scale shown). For each time point, a higher magnification of the boxed area is shown on the right. (**D**) FLIM value expressed as Flipper average lifetime (in ns) in the Golgi-positive area (delineated by the dashed white lines, zoom in images) was quantified at the indicated time points. Statistical analysis was performed using a mixed-effects model (REML) allowing for missing data points assuming sphericity and matching across biological replicates, correcting for multiple comparisons using Dunnett’s test (n≥3 replicates; N≥10 cells per replicate). (**E**) Images of live RPE1-ManII-Halo stably expressing cells plated on FN-coated glass coverslips, or stiff (30 kPa) or soft (2 kPa) FN-coated PAA gels. Flipper lifetime was measured as in (**C**). The Golgi apparatus was post-labelled using JF-646 (gray scale). (**F**) Flipper average lifetime in the Golgi-positive area was quantified and shown for the different conditions. Repeated-measures 1-way ANOVA tests were performed using Tukey’s post-hoc multiple comparison test (n=3 replicates; N∼30 cells per replicate). All scale bars, 30 µm.

We then examined the effect of two extracellular mechanical perturbations on Golgi tension. First, RPE1-ManII-Halo cells were seeded on FN-coated glass slides and allowed to spread for 0.5, 1, 2, 3, or 4 h. At each time point, cells were post-labeled with JF-646 Halo ligand to mark the Golgi area and Halo-Flipper, which was imaged by FLIM (***Fig. 3C***). Over the spreading time course, Halo-Flipper fluorescence lifetimes at the Golgi increased, suggesting a progressive rise in Golgi membrane tension that parallels FA assembly and cell spreading (***Fig. 3C, D***). Second, RPE1-ManII-Halo-expressing cells were seeded for 4h on FN-coated PAA gels of different stiffness or on glass (***Fig. 3E***). FLIM measurements showed reduced lifetimes on soft substrates compared with glass (***Fig. 3E, F***), suggesting reduced Golgi membrane tension under low mechanical load. However, while gradual differences between soft and intermediate-stiffness substrates, and between intermediate-stiffness substrates and glass were observed, they were more subtle and did not reach statistical significance in our analysis. Together, these data suggest that Golgi membrane mechanics is sensitive to extracellular mechanical cues, thereby linking adhesion-dependent cell states, cell spreading area, and post-Golgi carrier formation: robust spreading on FN-coated glass correlates with elevated Golgi tension and increased number of post-Golgi carriers, whereas limited adhesion on soft substrates corresponds to lower Golgi tension and reduced post-Golgi carrier abundance. These findings suggest that Golgi membrane mechanics responds to external mechanical forces, pointing to a potential mechanoadaptive regulation of Golgi function.

### Mechanical cues drive MT acetylation to enhance Golgi export

We next investigated how Golgi membranes respond to mechanical cues at the cell surface and what signaling pathways may relay forces to the Golgi apparatus. It is known that extracellular mechanical signals tune tubulin acetylation and GEF-H1/RhoA activation (Seetharaman et al., 2022; Krendel et al., 2002). GEF-H1 is a MT-associated RhoA exchange factor, which, upon MT acetylation, is released from MTs and is able to activate RhoA, providing a potential link between MT acetylation and downstream RhoA signaling (Krendel et al., 2002; Deb Roy et al., 2026). GEF-H1/RhoA activation can in turn lead to PLCε activation and DAG production at the TGN (Eisler et al., 2018). DAG is necessary for CARTS biogenesis by recruiting and activating PKD, a kinase required for the fission of Golgi-to-PM carriers such as CARTS (Wakana and Campelo, 2021). Based on this body of evidence, we hypothesized that MT acetylation could serve as one of the mediators transmitting mechanical signals from the cell surface to the Golgi membranes. Because astrocytes plated on stiffer substrates show higher levels of acetylated MTs (Seetharaman et al., 2022), we tested whether other mechanical stimuli –cell spreading on different ligand coatings and equibiaxial cell stretch– similarly induce MT acetylation.

First, to test whether MT acetylation can mediate mechanotransduction from FAs to the Golgi membranes, we compared MT acetylation levels in HeLa cells seeded on FN to those in cells seeded on PLL (***Fig. 4A***). By 1 h on FN, acetylated tubulin appears to be mostly localized in the perinuclear area (***Fig. 4A***). By 4 h, long protruding acetylated MTs extended from the perinuclear region toward the cell periphery (***Fig. 4A***). This pattern was absent in PLL-plated cells at all time points (***Fig. 4A***). Quantification of the acetylated-to-total tubulin ratio revealed a time-dependent increase on FN that plateaued at 4 h, whereas cells on PLL displayed no consistent trend, possibly reflecting large cell-to-cell variability and lack of integrin engagement on PLL substrates (***Fig. 4B***). Because changes in MT acetylation can modulate GEF-H1/Rho signaling, we next asked whether this spreading-dependent increase in MT acetylation was accompanied by activation of the downstream Rho/ROCK-Myosin II contractility axis. Rho activation leads to ROCK activation, which promotes Myosin light chain (MLC) phosphorylation on Ser19 (Totsukawa et al., 2000). In our previous work (Miserey-Lenkei et al., 2017, 2010), we showed that phosphorylated MLC (pMLC) associates with RAB6-positive carriers at the Golgi and that Myosin II activity is required for the biogenesis of RAB6-positive post-Golgi carriers at the TGN membrane. We therefore assessed MLC phosphorylation at Ser19 as a reporter of Myosin II activation downstream of Rho/ROCK signaling. Quantification at the basal cell-substrate interface revealed that pMLC levels increased at the earliest time point examined (1h) after seeding on FN, but not on PLL, and then returned toward steady-state levels at 4h and 24h (***Fig. S3A, B***). In parallel, F-actin staining using phalloidin confirmed the formation of stress fibers, another hallmark of Rho/ROCK signaling, in cells seeded on FN, in contrast to cells seeded on PLL (***Fig. S3A***). Although additional experiments will be needed to fully establish the significance of the transient increase in pMLC levels, these results are consistent with an enhanced Rho-dependent signaling at early stages of cell spreading on FN.

**Figure 4.**
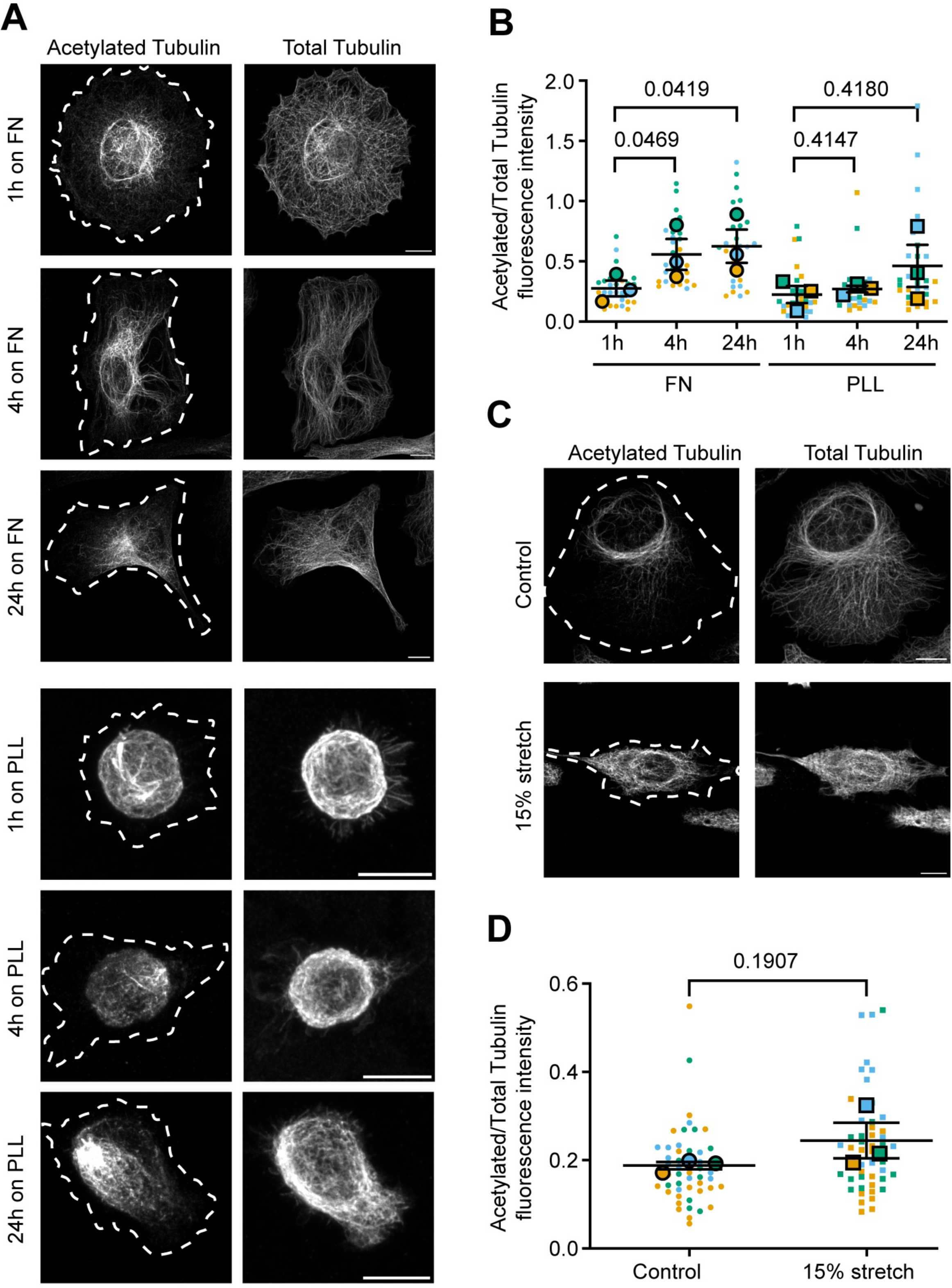
Cell spreading induces tubulin acetylation. (**A**) Representative confocal microscopy images of the acetylated and total α-tubulin fractions of fixed HeLa cells seeded for the indicated times on FN- or PLL-coated glass coverslips. Scale bars, 10 µm. Cell contours are indicated by dashed white curves. (**B**) SuperPlot showing individual cell measurements (small, light-colored symbols; N∼10 per biological replicate) and the mean value for each independent biological replicate (larger, black-outlined circles; n=3 replicates). Each color represents a different experimental replicate. A repeated-measures 2-way ANOVA test was performed for FN-seeded cells. P values using Fisher’s LSD test are reported in the figure. (**C**) Representative confocal microscopy images of the acetylated and total α-tubulin fractions of fixed HeLa cells seeded over FN-coated PDMS membranes, and subjected to no external forces of 15% mechanical strain. Scale bars, 10 µm. (**D**) SuperPlot showing individual cell measurements (small, light-colored symbols; N∼10 per biological replicate) and the mean value for each independent biological replicate (larger, black-outlined circles; n=3). Each color represents a different experimental replicate. A two-sided parametric ratio paired t-test was used. The P value is indicated in the plot.

Next, to emulate externally applied stretch, we subjected HeLa cells to a 15% equibiaxial strain for 30 min using our stretching device (***Fig. 2A***). External stretch suggested a trend toward higher tubulin acetylation relative to unstretched controls (***Fig. 4C***), but the effect was heterogeneous across experiments and, overall, not statistically significant, indicating that additional factors may modulate this response (***Fig. 4D***). This is consistent with recent work showing that cyclic mechanical stimulation can reorganize acetylated MT architecture without necessarily increasing total acetylated MT levels, highlighting that different mechanical cues may affect MT acetylation and organization in distinct ways (Li et al., 2023).

To determine whether MT acetylation alone is sufficient to increase the number of Golgi-derived carriers independently of external mechanical forces, we treated HeLa cells with tubacin, a histone deacetylase family member 6 (HDAC6) inhibitor that increases MT acetylation. As reported previously (Haggarty et al., 2003a; b), tubacin elevated acetylated tubulin levels (***Fig. S4A, B***), and significantly increased the number of PAUF-mRFP–positive carriers released from the TGN, without any significant increase in adhesion area (***Fig. 5A-D, Fig. S4C***). Tubacin treatment in RPE1-ManII-Halo-expressing cells did not lead to any significant increase in Golgi membrane tension as measured by Halo-Flipper FLIM (***Fig. 5E,F***), suggesting that signaling pathways other than those downstream of tubulin acetylation may control Golgi membrane tension and function. Nevertheless, because HDAC6 has substrates beyond tubulin, we cannot exclude additional, indirect effects of tubacin on adhesion- or cytoskeleton-dependent signaling. Collectively, these results indicate that physiological spreading on FN enhances MT acetylation, whereas acute, sustained stretch produces only a variable trend in the same direction. In addition, pharmacological induction of MT hyperacetylation with tubacin is sufficient to promote Golgi export. Thus, MT acetylation emerges as an important component of the mechanotransduction pathway from the cell surface to the Golgi apparatus, while our data suggest that additional routes likely contribute to the regulation of Golgi membrane tension and function. Future experiments using hypo- and hyper-acetylated MT expression constructs (Joo and Yamada, 2014) may provide a more direct means of defining the precise role of MT acetylation and how this is integrated with parallel pathways to coordinate Golgi mechanoresponse and mechanosensitivity.

**Figure 5.**
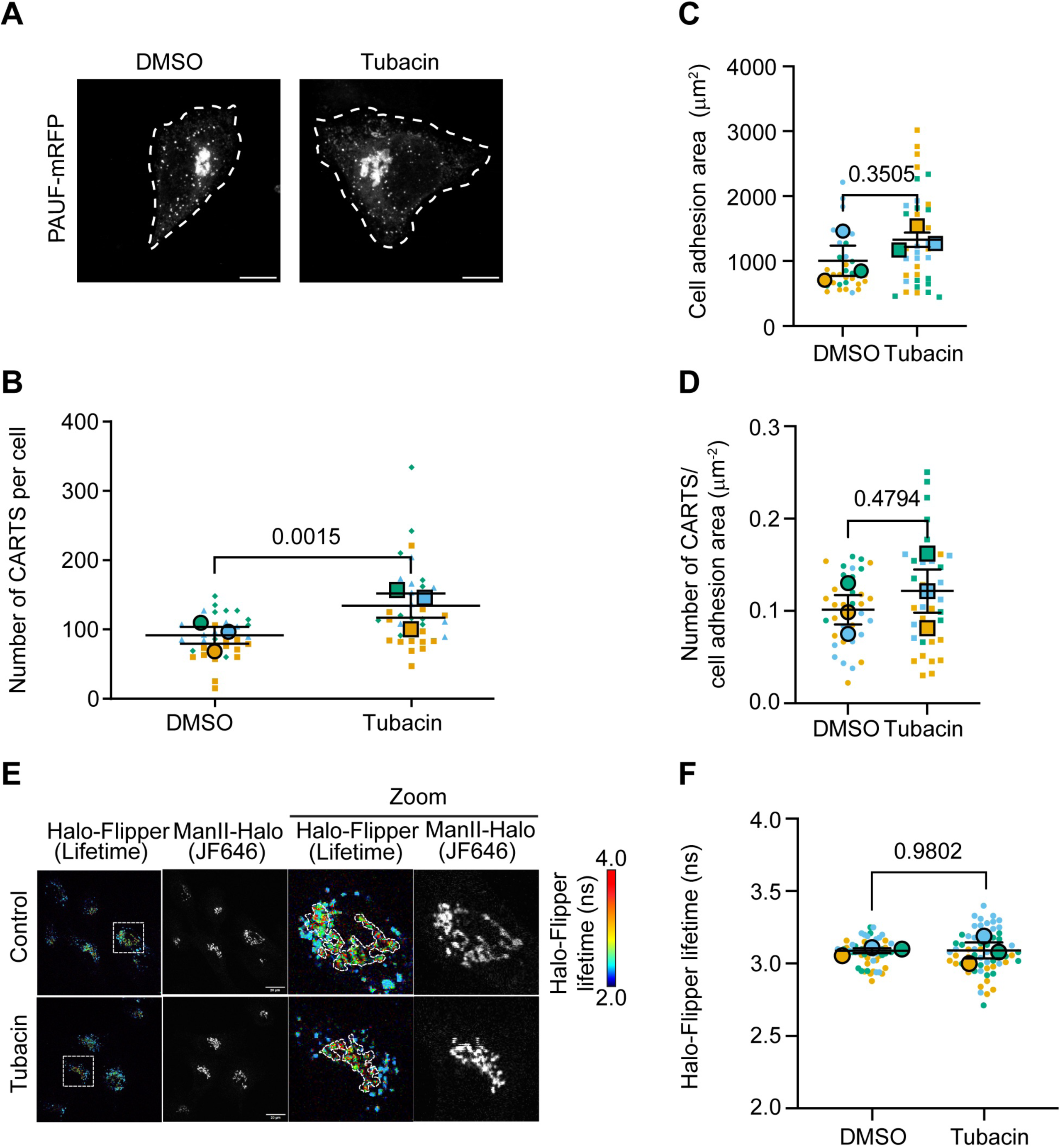
MT hyperacetylation promotes CARTS biogenesis. (**A**) Confocal microscopy images of fixed HeLa cells transfected with the PAUF-RFP plasmid, subjected to a treatment with or without tubacin. Cell contours are indicated by dashed white curves. Scale bars, 10 µm. (**B-D**) SuperPlots showing individual cell measurements (small, light-colored symbols; N∼10 per biological replicate) and the mean value for each independent biological replicate (larger, black-outlined circles; n=3) of (**B**) number of CARTS per cell, **(C)** cell adhesion area, and **(D)** number of CARTS per cell adhesion area. (**E**) FLIM Images of live RPE1-ManII-Halo stably expressing cells before and after overnight treatment with 10 µM of tubacin. The FLIM signal (Halo-Flipper lifetime) is displayed. Golgi was post-labelled using JF-646 (intensity shown, JF646 channel). Higher magnification of the boxed areas is shown on the right. Scale bar, 20 µm. (**F**) FLIM-measured Halo-Flipper fluorescence lifetime (in ns) was quantified for each experimental condition (n=3 replicates; N∼10 cells per replicate). Two-sided parametric ratio paired t-tests were used in all plots and P values are indicated in the plots.

### Golgi DAG levels and PKD activity are modulated by mechanical stimuli

Having shown that mechanical cues and MT acetylation enhance CARTS formation, we next asked whether these inputs also regulate Golgi DAG levels and PKD activity, since DAG recruits and activates PKD to promote carrier biogenesis downstream of RhoA/PLCε signaling (Eisler et al., 2018). To monitor DAG at the Golgi area, HeLa cells were transfected with the GST-C1a-PKD DAG biosensor (Maeda et al., 2001) and analyzed by immunofluorescence microscopy. We note that, as with other lipid probes based on functional lipid-binding domains, this sensor has limitations, including the possibility that it may not fully capture transient DAG dynamics and could potentially interfere with endogenous protein-lipid interactions. First, we compared cells seeded on FN with cells seeded on PLL (***Fig. 6A***). Cells on FN accumulated higher Golgi DAG levels than cells on PLL, based on the GST-C1a-PKD DAG biosensor (***Fig. 6A, B***). In FN-seeded cells, Golgi DAG showed an increase at early time points, followed by a return toward steady-state levels at later time points. This may reflect rapid DAG turnover during transport carrier biogenesis, including incorporation into nascent carriers or conversion into other lipids involved in membrane fission (Fugmann et al., 2007; Campelo and Malhotra, 2012). By contrast, cells on PLL showed reduced levels of Golgi DAG at earlier time points as compared to 24 h. Second, we compared DAG levels at the Golgi area in cells on soft (2 kPa) versus stiff (30 kPa or glass) FN-coated PAA gels (***Fig. 6C, D***). Consistent with our spreading data, and paralleling increased MT acetylation and Golgi-derived carrier biogenesis under these conditions, cells on stiff substrates tended to show higher DAG content at the Golgi area than cells on soft substrates, although only the difference between 2 kPa and glass was statistically significant (***Fig. 6D***). Finally, to test whether MT acetylation alone modulates Golgi DAG levels, we treated HeLa cells with tubacin (***Fig. 6E, F***). Whereas tubacin-induced MT hyperacetylation increased the number of cytoplasmic CARTS (***Fig. 5B***), it caused only a modest, non-significant increase in Golgi DAG content (***Fig. 6E, F***). Consistently, Golgi membrane tension remained unchanged upon tubacin treatment (***Fig. 5E, F***), suggesting that full DAG accumulation and membrane tension increase at the Golgi membranes likely require additional force-dependent activation of other DAG-synthesizing pathways (Romani et al., 2019).

**Figure 6.**
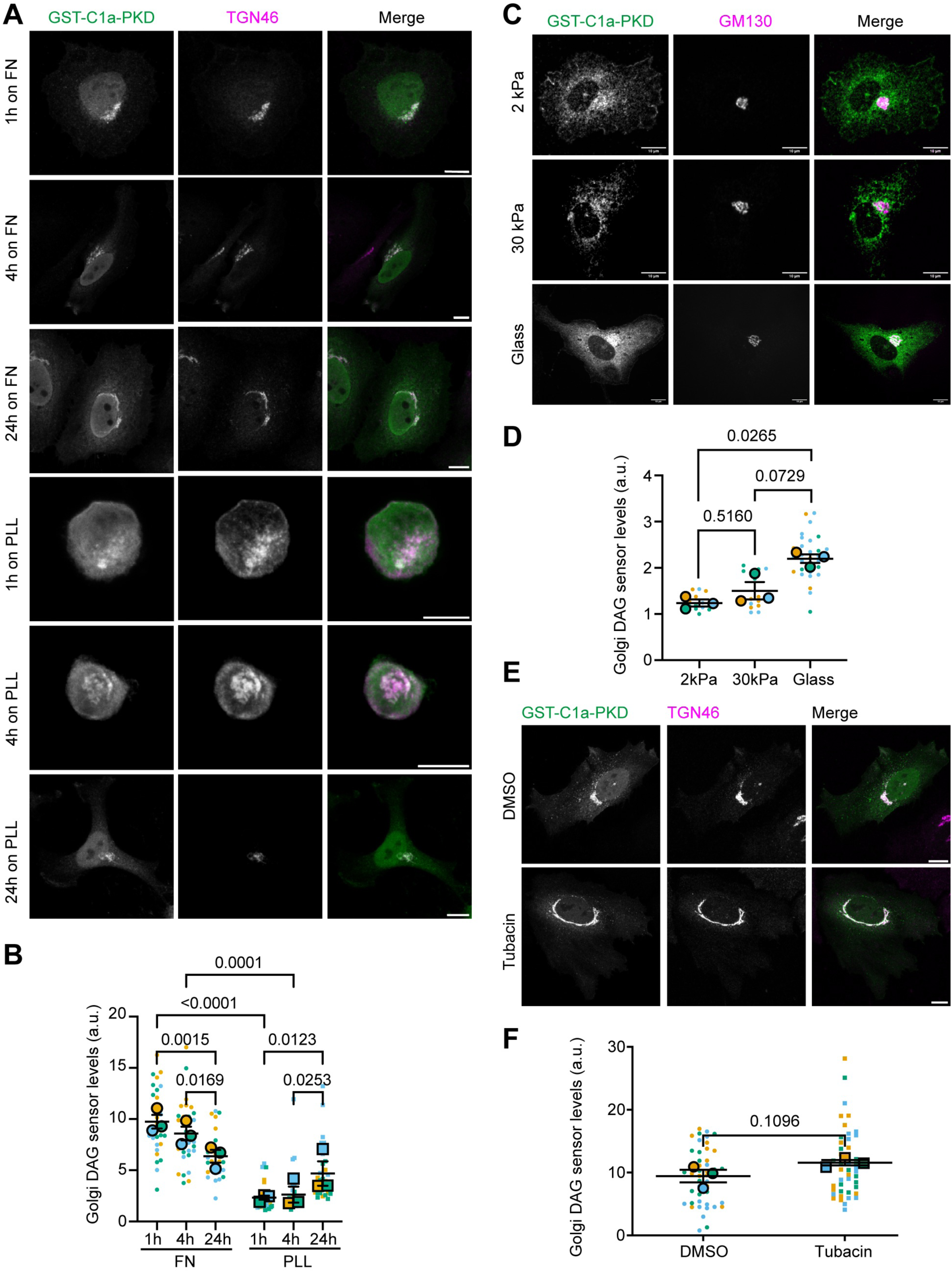
Mechanical cues alter Golgi DAG content. (**A**) Representative confocal microscopy images of HeLa cells transfected with the GST-C1a-PKD DAG sensor, seeded over FN or PLL for the indicated times, fixed, and immunostained for GST and TGN46. Scale bars, 10 µm. (**B**) SuperPlot (N∼10 cells per biological replicate; n=3 biological replicates) of the quantification of (**A**). A repeated-measures 2-way ANOVA test was performed. P values using Tukey’s post-hoc multiple comparison test are reported. (**C**) Representative images of HeLa cells transfected with the GST-C1a-PKD DAG sensor, seeded on FN-coated glass coverslips, stiff (30 kPa) or soft (2 kPa) PAA gels, fixed, and immunostained for GST and GM130. (**D**) SuperPlot showing quantification of (**C**). A repeated-measures 1-way ANOVA test was performed using Tukey’s post-hoc multiple comparison test (n=3 replicates; N≥5 cells per replicate). (**E**) Representative confocal microscopy images of HeLa cells transfected with the GST-C1a-PKD plasmid, subjected to a treatment with 10 µM Tubacin (or DMSO), fixed, and immunostained for GST and TGN46. Scale bars, 10 µm. (**F**) SuperPlot (N∼10 per biological replicate; n=3 biological replicates) showing quantification of (**E**). A two-sided parametric ratio paired t-test was used. The P value is indicated in the plot.

To complement these measurements, we directly monitored PKD activity at the Golgi/TGN using a genetically encoded reporter (G-PKDrep, see Methods, (Fuchs et al., 2009; Eisler et al., 2018)). PKD activity was higher at early time points after cell seeding on FN (***Fig. 7A, B***), consistent with the dynamics of DAG levels detected with the C1a-PKD-based biosensor (***Fig. 6B***) as well as of pMLC signal (***Fig. S3B***). Interestingly, higher Golgi PKD activity was also observed at early time points in cells seeded on PLL, indicating that the relationship between DAG/PKD signaling and Golgi export is more complex than captured by DAG measurements alone. While new tools capable of locally and quantitatively reporting lipid levels with sufficient sensitivity will be invaluable to validate and further dissect the dynamics of this lipid-mediated regulation of Golgi export, our findings suggest that DAG and PKD activity at the Golgi may be modulated by mechanical inputs, potentially as components of a mechanotransduction pathway linking cell surface mechanical forces to Golgi function.

**Figure 7.**
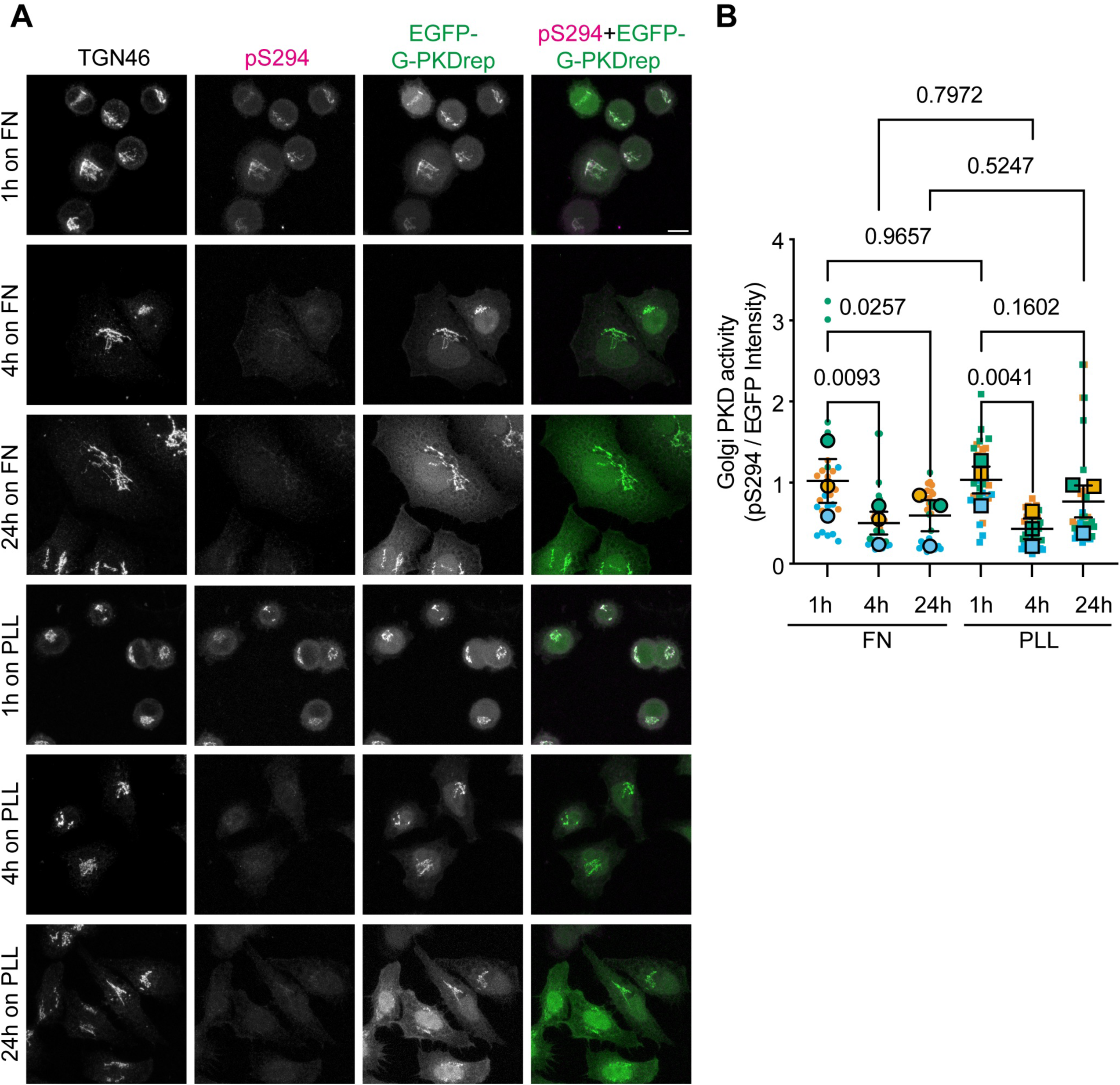
Mechanical cues modulate PKD activity at the Golgi membranes. (**A**) Representative confocal microscopy images of HeLa cells stably expressing the EGFP-tagged Golgi-localized PKD activity reporter (EGFP-G-PKDrep), seeded on FN or PLL for the indicated times, fixed, and immunostained for TGN46 and phosphorylated PKD substrate motif Ser294 (pS294). Scale bar, 10 µm. (B) SuperPlot (N∼10 cells per biological replicate; n=3 biological replicates) of the quantification of PKD activity, measured as the ratio between pS294 intensity and total EGFP-G-PKD signal at the Golgi. A repeated-measures 2-way ANOVA test was performed. P values using Tukey’s post-hoc multiple comparison test are reported.

### Golgi-derived export is necessary for cell spreading and mechanoadaptation

Our data so far point to a critical feedback loop between the cell surface –particularly FAs–and the Golgi apparatus, through which extracellular mechanical cues reshape Golgi lipid composition, signaling activity, mechanics, and secretory output. We therefore tested whether this feedback loop is functionally required for efficient cell adhesion and spreading. To investigate this, we performed spreading assays in HeLa cells expressing the membrane marker GFP-mem, while selectively impairing Golgi export using different strategies. First, we treated cells with Golgicide A, a general inhibitor of Golgi-to-PM trafficking (Saenz et al., 2009). Although initial cell spreading within the first hour was unaffected, a significant reduction in cell adhesion area was evident by 4h in Golgicide A-treated cells (***Fig. S5A, B***), suggesting that Golgi-derived export becomes an important contributor to continued spreading. We next inhibited PKD using CRT0066101 (Harikumar et al., 2010). PKD inhibition decreased the number of cytoplasmic CARTS (***Fig. 8A, B***) (Sugiura et al., 2026) and, similarly to Golgicide A treatment, led to a reduced expansion of the adhesion area at 4h (***Fig. 8C, D***). In parallel, FLIM measurements in RPE1-ManII-Halo cells showed that CRT0066101 treatment also reduced Golgi membrane tension, as reported by Flipper lifetime (***Fig. 8E, F***). To begin addressing how PKD inhibition affects FA organization, we examined active β1 integrin and FA area in cells treated with CRT0066101 during spreading on FN (***Fig. S5C***). PKD inhibition did not substantially change active β1 integrin levels at the basal cell surface at either 1h or 4h after seeding (***Fig. S5D***), whereas FA area was reduced after 4h (***Fig. S5E***), in parallel with the decrease in overall cell spreading area (***Fig. 8D***). These results suggest that inhibition of Golgi exocytic activity limits sustained cell spreading and is accompanied by changes in FA organization. Consistent with previous work showing that PKD2/PKD3 depletion does not significantly alter FA number per cell (Eisler et al., 2018), our data suggest that PKD-dependent Golgi export primarily supports sustained spreading downstream of adhesion-mediated signaling. This interpretation is also consistent with previous work showing that PKD activity supports directional migration and invasion in breast cancer models (Borges et al., 2015). Together, these findings indicate that disruption of PKD-dependent carrier biogenesis reduces both Golgi membrane tension and cell spreading efficiency. Under the same conditions, PKD inhibition did not significantly alter global MT acetylation levels after 4 h of spreading on FN (***Fig. S5F, G***), consistent with the idea that MT acetylation acts upstream of PKD activation. This also suggests that any feedback from PKD-dependent CARTS biogenesis onto MT acetylation is limited, at least at the level of global acetylated tubulin detected in our assay.

**Figure 8:**
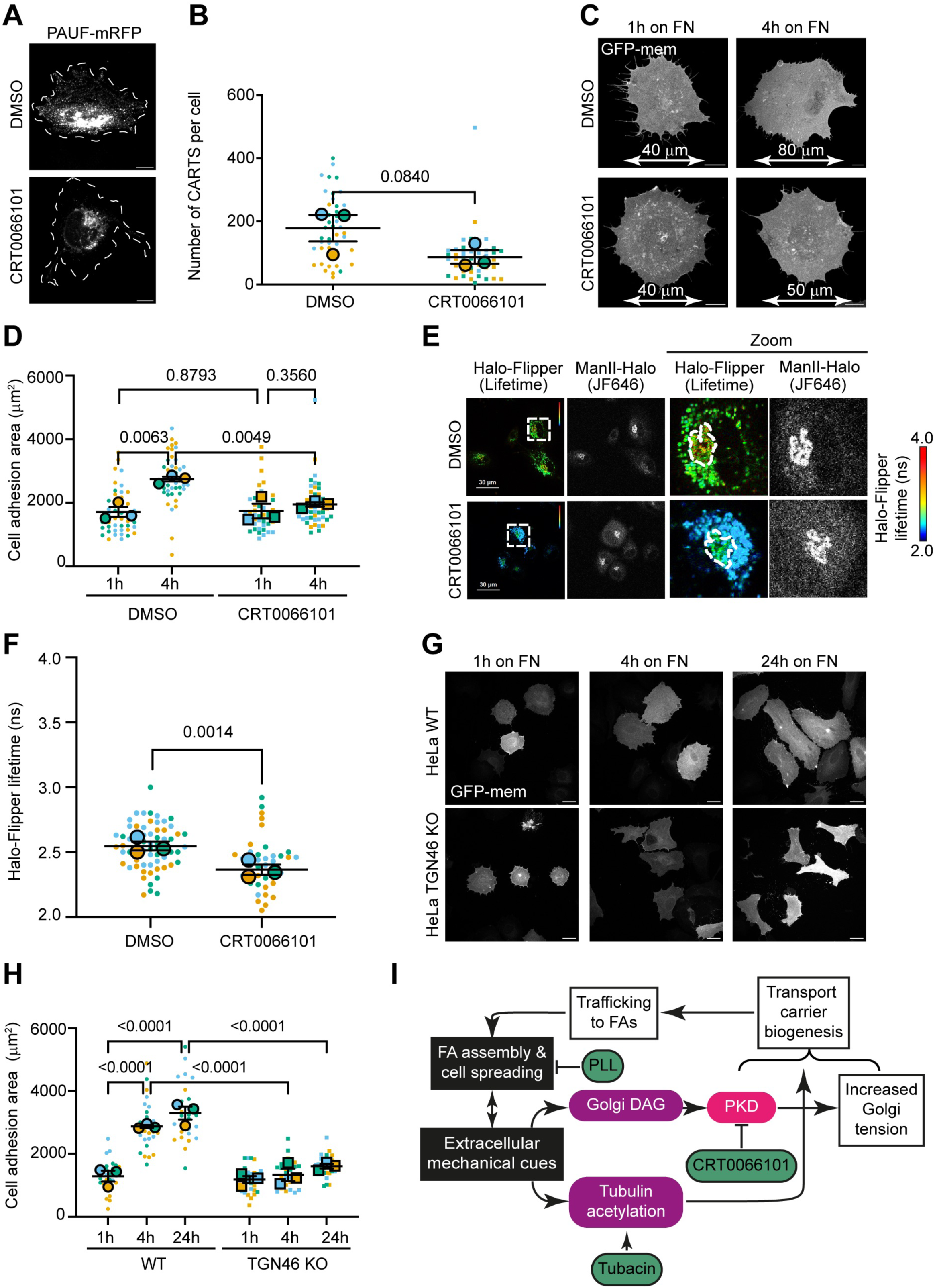
Golgi-derived export is necessary for cell spreading and mechanoadaptation. (**A**) HeLa cells transfected with the PAUF-RFP plasmid were spread on FN for 4 h and subjected to a treatment with (5 µM) or without CRT0066101, a PKD inhibitor, fixed, and imaged by confocal microscopy. Scale bars, 10 µm. **(B)** SuperPlot showing the number of CARTS per cell for each experimental condition. A two-sided parametric ratio paired t-test was used (n=3 replicates; N∼10 cells per replicate). **(C)** Representative confocal microscopy images of fixed HeLa cells transfected with the GFP-mem plasmid, subjected or not to a treatment with 5 µM CRT0066101. Scale bars, 10 µm; representative cell length scales shown. **(D)** SuperPlots showing individual cell measurements (N∼10 per biological replicate; n=3 biological replicates). A repeated-measures 2-way ANOVA test using Fisher’s LSD test was performed. (**E**) Images of live RPE1-ManII-Halo stably expressing cells before and after treatment with 5 µM CRT0066101. The FLIM signal and lifetime are displayed. The Golgi apparatus was post-labelled using JF-646. Scale bars, 30 µm. Higher magnification of the boxed areas is shown on the right. **(F)** SuperPlot showing Halo-Flipper lifetime values in the Golgi-positive area for each experimental condition (n=3 replicates; N∼10 cells per replicate). A two-sided parametric ratio paired t-test was used. The P value is indicated in the plot. **(G)** Representative confocal microscopy images of fixed HeLa cells (WT or TGN46 KO), transfected with the GFP-mem plasmid, and seeded over FN for the indicated times. Scale bars, 10µm. **(H)** SuperPlot showing individual cell measurements (N∼10 per biological replicate; n=3 biological replicates) quantifying adhesion area from images in (**G**). A repeated-measures 2-way ANOVA test was performed. P values using Tukey’s post-hoc multiple comparison test are reported. (**I**) Schematic representation of our findings (see text for details).

Because PKD also regulates actin dynamics and transcription (Fu and Rubin, 2011), we took a more targeted approach by selectively impairing CARTS-mediated export. HeLa TGN46 KO cells, which lack the transmembrane cargo adaptor TGN46 that is required for cargo sorting into CARTS and show impaired CARTS-mediated export (Lujan et al., 2024), exhibited significantly reduced spreading compared to wild-type (WT) cells, a phenotype that persisted for at least 24h post-seeding (***Fig. 8G, H***). Collectively, our results underscore that the Golgi apparatus actively responds and adapts to extracellular mechanical cues (***Fig. 8I***). The ability of the Golgi apparatus to sense, respond, and secrete in response to extracellular mechanical forces supports proper cell adhesion, spreading, and homeostasis. Our work unveils new concepts in organelle mechanobiology, highlighting key mechanotransduction steps at the Golgi apparatus, a secretory mechanoresponse, and the importance of feedback communication between the PM and internal membranes in maintaining cellular homeostasis.

## DISCUSSION

Our work reveals a mechanotransduction axis linking extracellular forces to the Golgi complex. These forces are transduced to remodel Golgi membranes by tuning their biophysical and biochemical characteristics, and lead to an increase in the number of cytoplasmic Golgi-derived carriers (CARTS as well as CD59-positive carriers) to support adhesion and efficient spreading. We show that three distinct mechanical stimuli (spreading forces on integrin-activating substrates, substrate stiffness, and equibiaxial stretch) associate with an increase in the number of Golgi-derived carriers. In parallel, several of these cues are accompanied with changes in Golgi membrane tension (measured by Halo-Flipper FLIM) and Golgi DAG/PKD signaling (measured by a fluorescent DAG biosensor and a PKD activity reporter). We provide evidence that MT acetylation acts as a critical downstream effector of mechanical cues and is sufficient to boost post-Golgi carrier formation even in the absence of external forces. Finally, we show that Golgi export –particularly via PKD- and CARTS-dependent routes– is both upregulated by mechanical stimuli and required for sustained cell spreading, suggesting a reciprocal communication between FA establishment, cell spreading, and directed Golgi-to-PM trafficking to exocytic hotspots.

Ample evidence supports the idea that exocytosis in adherent cells is not uniformly distributed across the cell surface but occurs at secretion hotspots, typically juxtaposed to FAs (Stehbens et al., 2014; Eisler et al., 2018; Huet-Calderwood et al., 2017; Fourriere et al., 2019). FAs are molecular platforms required for adhesion and for sensing and transducing extracellular mechanical forces (Kechagia et al., 2019; Chastney et al., 2025). Integrin β1 itself is delivered from the TGN to the PM through a RAB6-mediated (Huet-Calderwood et al., 2023) and PKD-dependent pathway (Yeaman et al., 2004), directly linking Golgi export routes to the reinforcement of adhesion complexes. Notably, recent work further showed that a subset of newly synthesized α_5_integrin integrins can be delivered rapidly and locally to adhesion-proximal plasma membrane regions through a Golgi-bypass route (Lerche et al., 2026). Together, these findings highlight that biosynthetic integrin trafficking is spatially regulated and can contribute directly to adhesion dynamics. However, whether –and how–these mechanical cues drive adaptation of the secretory machinery has remained largely unexplored. Notably, recent studies show that ER exit sites (Farhan et al., 2025) and ER export can be upregulated upon extracellular mechanical cues by the GTPase Rac1 (Phuyal and Baschieri, 2020; Phuyal et al., 2022), and optogenetic and other mechanostimulatory perturbations alter ER export dynamics (Song et al., 2024; Chen et al., 2025). In addition, FAs and ECM can regulate the COPII machinery involved in ER to Golgi transport (Jung et al., 2022). Collectively, these reports point to coordinated, adhesion-dependent regulation across the secretory pathway. In line with this emerging view, a study published during revision of this manuscript showed that substrate stiffness regulates post-Golgi cargo sorting and secretion through a Src-FAK-AMPK-GBF1 pathway, further supporting the idea that the Golgi functions as a mechanically regulated hub within the secretory pathway (Serafino et al., 2026). Our findings therefore position the Golgi as an active mechanoresponsive organelle, extending mechanotransduction beyond conventional mechanosensor organelles, such as the PM and nucleus.

Our data place MT acetylation downstream of extracellular mechanical inputs and upstream of Golgi functional adaptation, consistent with its role as a tension-sensitive mediator (Seetharaman et al., 2022) and a regulator of PKD-dependent Golgi trafficking (Eisler et al., 2018). By showing that inhibiting tubulin deacetylation (using the HDAC6 inhibitor tubacin) partially phenocopies mechanical stimulation, we identify MT acetylation as both a sensor and an amplifier of force signals, upstream to the GEF-H1/RhoA/PLCε/DAG/PKD axis previously implicated in Golgi export (Eisler et al., 2018). We propose a model (***Fig. 8I***) in which mechanical forces –transduced via a yet-unidentified mechanoreceptor– may trigger increased MT acetylation, release of GEF-H1, and RhoA signaling (Seetharaman et al., 2022). Active RhoA reinforces actomyosin contractility through ROCK and also promotes DAG production at the TGN via PLCε, recruiting and activating PKD to drive carrier biogenesis (Eisler et al., 2018) (***Fig. 8I***). Direct perturbation of GEF-H1 or alpha-tubulin acetylatransferase 1 (αTAT1), as well as the use of hypo- and hyper-acetylated tubulin constructs, will be important to further dissect the role of MT acetylation in this pathway. Too low Golgi membrane tension may render the TGN inefficient for carrier fission or alternatively may be a consequence of an altered TGN lipidome, so the full extent of this mechanical adaptation warrants further study. Of note, elevated membrane tension is known to work against membrane curvature generation (Dai and Sheetz, 1995; Carlsson, 2018; Le Roux et al., 2021), highlighting the importance of a well-balanced, dynamic control of Golgi membrane tension for secretory function. Importantly, our observations can be placed in the context of the opposing effects that membrane tension plays on endocytic versus exocytic pathways. High plasma-membrane tension is well-documented to inhibit endocytosis (Apodaca, 2002; Kosmalska et al., 2015; Wu et al., 2017), a block that can be overcome locally by polymerization of actin to provide the force required for vesicle invagination and scission (Boulant et al., 2011; Gauthier et al., 2011). By analogy, actomyosin contractility may not be purely inhibitory but also contribute to promote the biogenesis and fission of post-Golgi carriers (Miserey-Lenkei et al., 2010) by shaping Golgi membrane mechanics and promoting DAG/PKD-dependent scission; thus, actin dynamics can act as a context-dependent permissive or driving force for membrane trafficking depending on the compartment and the direction of membrane flow.

Notably, we found that tubacin-induced MT hyperacetylation induced only a modest increase in Golgi DAG and no measurable change in Golgi tension, suggesting that additional regulatory layers and other effectors, such as Lipin-1 (Romani et al., 2019), may contribute to full DAG accumulation, PKD activation, and/or tension modulation. We note, however, that Flipper probes report membrane lipid packing/ordering as a proxy for membrane tension, and perturbations that alter Golgi lipid composition (e.g., PKD inhibition or DAG modulation) can alter Flipper lifetimes independently of tension reporters (Colom et al., 2018). Thus, while our FLIM data are consistent with tension changes, interpretation of lifetime shifts as pure mechanical readouts should be cautious and will require complementary validation in future work.

Taken together, our data suggest that early cell spreading on FN triggers a transient Rho/ROCK-dependent program that couples actomyosin contractility to Golgi secretory activity, as reflected by the early increase in pMLC, DAG, and PKD activity and consistent with a role in rapid membrane remodeling and Golgi-derived carrier biogenesis. Previous work has shown that human fibroblasts display a faster and stronger increase in pMLC levels during spreading on FN as compared to PLL (Ren et al., 2004), consistent with differential activation of contractility pathways under these conditions. Using more specific activity reporters for Rho or GEF-H1, as well as tools such as optoTAT (Deb Roy et al., 2026), an optogenetic technology that allows for local induction of MT acetylation, will be instrumental to fully dissect the precise molecular links between MT acetylation, RhoA activation, Golgi membrane tension, and PKD activation –and, more broadly, the Golgi machinery that tunes secretory output (***Fig. 8I***). Nonetheless, by uncovering these connections, our work establishes Golgi-based mechanoadaptation as a key mechanism of cellular regulation and opens important avenues for future investigation.

Reciprocal feedback between exocytosis and membrane tension also provides a physiological rationale for the coupling we observed: exocytosis lowers PM tension by adding membrane facilitating further spreading and migration (Gauthier et al., 2011; Pontes et al., 2017). Thus, upregulation of Golgi-to-PM trafficking and secretion at adhesion-proximal hotspots can both respond to and modulate PM mechanics, promoting processes such as migration, tissue morphogenesis, and membrane homeostasis. In pathological contexts, such as cancer invasion and fibrosis, altered ECM mechanics could dysregulate this Golgi mechanoresponse, driving aberrant secretion of matrix components or signaling factors. Notably, delivery of newly synthesized integrin β1 relies on RAB6- and PKD-dependent Golgi export (Huet-Calderwood et al., 2017; Yeaman et al., 2004), suggesting that misregulation of this pathway could directly impact adhesion turnover and invasive behavior. Targeting elements of this axis (e.g., HDAC6, PLCε, PKD, TGN46, or other intermediates yet to be identified) may offer novel therapeutic strategies to modulate mechanoresponsive secretion. Future experiments using micropatterned substrates of defined size and geometry (Théry, 2010), ideally combined with substrates of tunable stiffness, will be important to separate the relative contributions of cell spreading area, FA organization, and mechanical load to Golgi carrier biogenesis and membrane tension regulation. Finally, extending these studies to polarized cells, 3D matrices, and *in vivo* models will be essential to understand Golgi-mediated mechanoregulation at the tissue level.

In summary, we uncovered a two-way dialogue between the PM and the Golgi apparatus that is mediated by mechanical forces, MT acetylation, and TGN signaling (DAG and PKD). This feedback loop may allow cells to adapt their adhesive and migratory behavior to the mechanical properties of their environment, thus positioning the Golgi as a new mode of cellular mechanoregulation.

## METHODS

### Reagents, plasmids, antibodies

The Paxillin-pEGFP plasmid was a gift from Rick Horwitz (Addgene plasmid #15233) (Laukaitis et al., 2001). The mKate2-FM4-PAUF plasmid was described earlier (Wakana et al., 2021). The Double palmitoylated Neuromodulin (N-terminal)-GFP (GFP-mem) plasmid is used as a general PM marker as it is a fusion protein with a signal for post-translational palmitoylation targeting the fusion protein to the cell membrane, and was a gift from Francesc Tebar’s lab (Universitat de Barcelona, Spain) (Vidal-Quadras et al., 2011). The plasmids encoding for PAUF-mRFP (Wakana et al., 2012) and GST-C1a-PKD (Maeda et al., 2001) were donated by Vivek Malhotra (CRG, Barcelona, Spain), and the plasmid encoding for PAUF-MycHis (Kim et al., 2009) by S. S. Koh (Korea Research Institute of Bioscience and Biotechnology, Daejeon, Korea). Commercial antibodies used in this study were as follows: sheep anti-TGN46 (Bio-Rad, AHP500GT); mouse anti-GM130 (BD Biosciences, 610822); mouse anti-paxillin (BD Biosciences, 610568); mouse anti-paxillin, clone 349 (BD Transduction Laboratories, 610051); rat anti-active β1 integrin, clone 9EG7 (BD Biosciences, 553715); rabbit anti-α-tubulin (Abcam, ab18251); rabbit anti-tRFP (Evrogen, AB233); mouse anti-acetylated α-tubulin (Sigma-Aldrich, T6793); rabbit anti-glutathione-S-transferase (GST) (Sigma-Aldrich, G7781); rabbit anti-ERC1/ELKS (Proteintech, 22211-1-AP); mouse anti-α-tubulin, clone B-5-1-2 (Sigma-Aldrich, T6074); mouse anti-Myc, clone 9E10 (Santa Cruz Biotechnology, sc-40); rabbit anti-Myc (Cell Signaling Technology, 2272); and mouse anti-pMLC Ser19 (Cell Signaling Technology, 3675). The pSer294-specific rabbit polyclonal antibody used for detection of G-PKDrep phosphorylation was kindly provided by Angelika Hausser (University of Stuttgart, Germany) and previously described (Hausser et al., 2005; Fuchs et al., 2009). Phalloidin-Alexa Fluor 647 was from Invitrogen and used at a 533-fold dilution from the stock (0.00375 U/µl final concentration) for 1 h. Secondary antibodies used were Alexa Fluor 488 (A32766), Alexa Fluor 555 (A32794), or Alexa Fluor 647 (A21448) coupled donkey anti-mouse, anti-rabbit and anti-sheep immunoglobulin G (IgG); as well as goat anti-mouse Alexa Fluor 488 (A11029), anti-rabbit Alexa Fluor 488 (A11034), and anti-mouse Alexa Fluor 594 (A11032) (all purchased from Invitrogen). Janelia Fluor probes (JF-646) were from Promega. The following reagents were purchased from Sigma-Aldrich: Fibronectin (11051407001), Poly-L-Lysine (PLL) (P1524), Tubacin (SML0065), Golgicide A (345862), and Cycloheximide (CHX) (C4859). D/D solubilizer (635054) was obtained from Takara Bio. Fibronectin (354008) was also obtained from Corning. CRT0066101 was from Tocris Bioscience (4975). Sylgard™ 184 Elastomer KIT was from Dow Inc (24001673921).

### Cell culture and transfection

HeLa cells were cultured in DMEM (Capricorn Scientific GmbH) supplemented with 10% FBS (Invitrogen), 1% penicillin-streptomycin (Gibco) and L-Glutamine (LabClinics) under 5% CO_2_ at 37°C. HeLa Cells were transiently transfected using X-tremeGENE™ 9 DNA transfection reagent (Sigma-Aldrich) following the manufactureŕs instructions. Cells were used for the designed experiments ∼16h post-transfection. In experiments where cells were placed at 20°C for 2h to synchronize cargo release from the Golgi apparatus, cell culture medium was supplemented with HEPES 25mM (Sigma-Aldrich). Additionally, cell culture medium was supplemented with CHX 100 µM to inhibit new protein synthesis before cargo release. TGN46 KO HeLa cells were generated as described previously (Lujan et al., 2024). RPE1 cells were cultured in DMEM/ F12 supplemented with 10% fetal bovine serum (FBS) under 5% CO_2_ at 37°C. RPE1 cells were transfected using Lipofectamine 3000 (Invitrogen) following the manufactureŕs instructions. RPE-ManII-Halo stable cell line was generated as follows: RPE1 cells were transfected with pIRES-ManII-Halo (ManII-Halo sequence from prHom-ManII-Halo, kind gift from D. Toomre, was inserted into pIRES-Neo3) and single clones were then selected following Ampicillin selection. HeLa Flp-IN T-Rex GPKDrep cell line (Hela-PKDrep) (Fuchs et al., 2009; Eisler et al., 2018) was kindly provided by Angelika Hausser (University of Stuttgart, Germany). HeLa cells stably expressing PAUF-MycHis were generated and described in (Sugiura et al., 2026).

### ELKS knockdown in HeLa cells stably expressing PAUF-MycHis

HeLa cells stably expressing PAUF-MycHis were transfected with control siRNA or an siRNA oligo targeting ELKS. The targeting sequences of siRNA were as follows: Control (GL2 luciferase): 5’-AACGTACGCGGAATACTTCGA-3’; ELKS: 5’-AAGGAAGTATTAAGAGAAAAT-3’. For immunofluorescence, at 48 h after siRNA transfection, the cells were trypsinized and plated on FN-coated coverslips; 24 h later, the cells were fixed with 4% PFA and analyzed by fluorescence microscopy. For testing knockdown efficiency, at 72 h after siRNA transfection, the cells were lysed with 0.5% SDS and 0.025 U/μl benzonase nuclease (Sigma-Aldrich) in PBS. The cell lysates were analyzed by Western blotting with anti-ELKS and anti-a-tubulin antibodies.

### Cell treatments

For MT acetylation and DAG production experiments, cells were pre-seeded for 1h on FN-coated coverslips (10µg/ml final concentration, overnight (o/n) incubation) and then treated with 10 µM Tubacin or DMSO for 3h continuously. For CARTS formation experiments, cells pre-seeded on FN-coated coverslips were subjected to a pre-treatment of 30 minutes with Tubacin or DMSO, followed by a 2h incubation at 20°C in presence of Tubacin or DMSO, and a 30 minutes CARTS release at 37°C with Tubacin or DMSO still present in the samples. For PKD inhibition experiments, cells were pre-seeded over FN-coated coverslips for 30 minutes and then incubated with CRT0066101 (5µM) or DMSO for 30 min or 210 min. For actin depolymerization, cells were labelled with Halo flipper and JF-646 as explained and then further treated with 50 nM latrunculin-A for 10 min and immediately imaged for the following 10–15 min. Imaging was discontinued as soon as Golgi fragmentation was observed in the cells, which was normally occurring at 25–30 min post-treatment. For Golgi export inhibition experiments, 30 min pre-seeded cells were treated with Golgicide A (10 µM) for 30 min or 210 min. The mKate2-FM4-PAUF construct contains the FM4 aggregation domain, retaining the cargoes in the ER after synthesis. By adding to the cell medium D/D solubilizer, a small molecule that solubilizes FM4-induced aggregates, it is possible to synchronize a wave of cargo release from the ER and follow their intracellular localization in time, all the way from the ER to the Golgi apparatus and to CARTS for secretion. For those cells transiently transfected with mKate2-FM4-PAUF plasmid, D/D solubilizer was added at 1µM final concentration to allow the cargoes to be exported out of the ER.

### Combined spreading-secretion assay

For cell spreading assays (e.g., ***Fig. 1A, B***), HeLa cells transiently transfected with GFP-mem and mKate2-FM4-PAUF or RPE1-ManII-Halo stable cells were split and seeded over a ligand-coated coverslip (FN or PLL). Unless otherwise stated, cells were lifted by incubation with Trypsin (0.05% trypsin, 0.53 mM EDTA) for 3–5 min in a 37°C incubator. When lifted using EDTA, cells were incubated with 10 mM EDTA in Ca^2+^ and Mg^2+^ free PBS for 10 min in a 37°C incubator. Cells were kept at 37°C, and 30 min before the end of the spreading assay, cell culture medium was supplemented with CHX (100 µM) and D/D solubilizer (1 µM) to allow cargoes to be released out of the ER. Cells were then fixed and visualized by fluorescence microscopy.

### Immunofluorescence and fluorescence microscopy imaging

Cells were fixed with 4% (v/v) paraformaldehyde (PFA) in PBS for 15 minutes at room temperature (RT). After three washes with PBS, cells were permeabilized with 0.2% Triton X-100 (TX-100) in PBS for 30 minutes at RT. After washing with PBS, blocking was performed with 4% bovine serum albumin (BSA) for 30 minutes at RT or o/n at 4°C. After washing the samples, the BSA solution at 2% was used to dilute the primary and secondary antibodies for incubation 1h at RT. Samples were then mounted on glass slides using either ProLong Gold Antifade Reagent (Thermo Fisher Scientific) or Mowiol. Widefield microscopy images were acquired on an upright Nikon Eclipse Ni-U microscope equipped with a 60x water-dipping objective (NIR Apo 60x/WD 2.8, Nikon), an ORCA-Flash 4.0 camera (Hamamatsu), and controlled by Metamorph software. Confocal microscopy images were acquired on a TCS SP8 microscope (Leica Microsystems GmbH, Germany) equipped with an HC PL APO CS2 100x/1.40 oil objective, a pulsed white light laser (WLL) operating at 20 MHz repetition rate, 70% master power and hybrid detectors (HyD) in photon-counting mode and 8-bit depth; whereas spinning disk confocal microscopy images were acquired on a Nikon Inverted Eclipse Ti-E (Nikon) microscope equipped with a Spinning disk CSU-X1 (Yokogawa), an iXon EMCCD camera (Andor) or sCMOS Kinetx 22 camera (Photometrics) integrated in Metamorph software by Gataca Systems, using 60x or 100x CFI plan apochromat VC 1.4 NA oil immersion objectives (Nikon). The spinning disk microscope was in a cage incubator from Life Imaging Services for temperature control. GFP, mKate2, RFP, Alexa488, Alexa647 were excited using 489, 589, 554, 499, or 653 nm lines, respectively, and the emission was detected between 499-580, 599-770, 564-750, 509-600, or 663-781 nm, respectively. For dual and triple-color imaging, sequential scan mode between lines was used to minimize crosstalk and mechanical drift. For z-stack acquisitions, the distance between confocal planes was set as system optimized by the microscope acquisition software, the planes were reaching across the entire cell volume, and maximum intensity (for visual representation) or sum slices (for total volumetric intensity measurements) projections were computed. Images were typically acquired with 2x line accumulation. Laser power was adjusted to avoid pixel saturation and then kept constant throughout each experiment. Image analysis was performed using ImageJ software.

### Selective protein immobilization (SPI) assay

The assay was firstly described in (Fourriere et al., 2019). Briefly, coverslips were incubated with bicarbonate buffer 100mM for 1h at 37°C. Then, coverslips were washed three times in PBS and dried before incubation with the anti-tRFP antibody for 3h at 37°C or o/n at 4°C, using a 1:250 dilution from the stock concentration. For the detection of the coated antibody to validate uniformity of coverslip coating, an anti-rabbit Star Orange secondary antibody (Abberior) was used.

### Live cell imaging

Images were acquired with a commercial Nikon Eclipse Ti System, equipped with a 100x oil objective with NA 1.49 using TIRF illumination and enclosed in an incubator chamber set to 37 °C and 5% CO_2_. The detection was carried out using an ANDOR technology EMCCD iXon 897 camera. The equipment presents an Agilent technologies laser box with wavelengths of 488 and 560 nm. Laser power was adjusted for each channel to prevent saturation. For the experiments tracking post-Golgi carriers transport to FAs, cells were seeded on FN-coated 35 mm glass bottom dishes (MatTek), and cells were subjected to a cargo release synchronization before imaging. Before cargo channel acquisition, an image of the FAs channel was taken as a reference. Then, cargo channel was acquired live at 1 fps for 45 min. For SPI live cell imaging, both channels were acquired quasi simultaneously every 30s for 1h. All live-cell imaging was acquired at 37°C.

### Retention using selective hooks (RUSH) assay

The RUSH assay was performed as described in (Boncompain et al., 2012).

### Measurement of PKD activity at the Golgi membranes

To measure PKD activity, we used a HeLa Flp-IN T-Rex GPKDrep cell line (Fuchs et al., 2009; Eisler et al., 2018) kindly provided by Angelika Hausser (University of Stuttgart, Germany), cultured under selection with 500 µg/ml Hygromycin B and 10 µg/ml Blasticidin. Prior to the experiment, antibiotics were removed and cells were induced using 10 ng/mL for 24h. The day after, cells in suspension were seeded on FN-coated coverslips (10 µg/mL) for the indicated times and processed for immunofluorescence. Image analysis pipeline consisted of segmenting the Golgi regions using the EGFP mask and measuring the total intensity of pS294 signal and EGFP signal for normalization. The ratio between pS294 and EGFP is indicative of Golgi PKD activity.

### Fluorescence lifetime imaging microscopy (FLIM)

A FLIM module from PicoQuant (Kit LSM Upgrade pour Nikon A1 PicoQuant, Opton Laser International) was added to the A1R Nikon confocal microscope with both resonant and galvano mode scanners. Images were acquired at 37°C, and by using 485 nm laser excitation and a 27.5ns dwell time per pixel with 5-frame acquisition. The emission bandpass filter wavelength was 506-619 nm. Single-stack images were captured and used for FLIM using TCSPC (Time-Correlated Single Photon Counting). Images were analyzed using the SymPhoTime 64 software (PicoQuant, Opton Laser International). The image acquisition time was typically 48 seconds for a 512×512 image. A region of interest was drawn manually based on a Golgi mask (Janelia Fluor® 646 HaloTag channel, see below for details) using the Screen Dragon software. The photon arrival time histograms were fitted with a double exponential model, after deconvolution with the calculated impulse response function in SymPhoTime. Upon fitting, the intensity weighted average fluorescence lifetime was computed and used for subsequent analysis. All FLIM was performing in living cells at 37°C.

### Halo-Flipper

ManII-Halo RPE1 cells were labeled with Halo-Flipper to report lipid packing defects in Golgi membranes due changes in membrane tension or lipid composition. Cells were plated on Fluorodishes with glass or PAA gel substrates from 30 min to 4h (depending on the experiment). Halo-Flipper (kindly provided by S. Matile Lab) of 100 nM concentration was prepared in 200 µl serum-free medium and added to the cells by replacing the old medium. After a 15 min incubation, Halo-Flipper was replaced with 160 µl complete medium (containing 10% FBS) with 40 µl of 1 µM Janelia Fluor® 646 HaloTag (JF-646, 200 nM final working concentration) and further incubated for 15 min, before FLIM and fluorescence images of the Golgi membrane were acquired. Halo-Flippers were gently imaged avoiding as much as possible light exposure of the samples before FLIM recording. To minimize possible Flipper-induced phototoxicity and singlet oxygen photosensitization (Torra et al., 2024), individual cells were measured only once. Halo-Flippers were made commercially available in 2025 and are now supplied by Spirochrome (Halo-Flipper, Spirochrome).

### Janelia Fluor HaloTag ligands

Janelia Fluor® 646 HaloTag (JF-646) was used at 0.5 µM (1:2000 dilution from the 1 mM stock) in culture media. Cells were labelled by incubation for 15 mins at 37 °C, followed by extensive washing.

### Preparation of polyacrylamide (PAA) gels

PAA gels were prepared as described in (Pérez-González et al., 2021).

### Preparation of PDMS stretchable membranes and ring mounting

The detailed protocol is described in (Le Roux et al., 2025). Briefly, the PDMS mixture was prepared in a 1:10 ratio and degassed in a vacuum chamber for 1 h. PDMS was spread over PMMA plates using a spin coater in 2 steps: a first step during 5 s at 500 rpm with a 100 rpm/s acceleration, and a second step during 1 min at 500 rpm with a 300 rpm/s acceleration. PDMS was polymerized o/n at 65°C in an oven. PDMS membranes were peeled off from the PMMA plates and mounted on the stretch-rings. PDMS rings were sterilized by exposing them to UV light for 15 min and then were coated with FN o/n. The following day, FN excess was rinsed off and cells were seeded on top of the FN-coated PDMS membranes for 1h prior the experiment. For experiments carried out with non-transiently transfected cells, cells were subjected to 15% substrate stretch for 30 min at 37°C before fixation. In experiments characterizing the formation of CARTS upon mechanical strain stimulation, an intermediate step at 20°C for 2h to synchronize cargo release was set before stretching. Control experiments were performed under the same conditions, but in the absence of mechanical strain. Cells subjected to mechanical forces were fixed and imaged under stretch to prevent visual aberrations.

### Quantification of images and movies

Analysis of the cell area was performed using a custom-designed macro in ImageJ. In brief, for each image cell area was determined by thresholding the GFP-mem channel, converting it to a mask and creating the selection. To determine number of CARTS, the ImageJ “Detect particles (ComDet)” plugin developed by Eugene A. Katrukha (Utrecht University, the Netherlands) was used, avoiding the perinuclear Golgi-positive area, using a maximum intensity z-projection of the entire confocal fluorescence microscopy z-stack of the cell or the epifluorescence image of the cell (when imaging the substrate stretching experiments). The normalized total intensity levels of acetylated tubulin were calculated according to the equation: Ratio of acetylated/total tubulin level = Acetylated tubulin intensity/Total tubulin intensity. DAG production (area and intensity) and Golgi area were calculated manually drawing an ROI around the fluorescent-positive mass present in the perinuclear area of the cells. In the bulk SPI analysis, FAs and secreted cargo objects were segmented and the localization and fluorescence intensity of each pixel was calculated. The nearest neighbor distance (nnd) between each pixel of secreted cargo to the closest pixel of FAs was measured using Matlab. The intensity of each pixel of secreted cargo was also included to obtain the 2D plots. CD59-positive vesicle count analysis was carried out using the cells expressing RUSH cargo of interest, which were fixed at 45 min post-biotin addition to record post-Golgi carriers. The vesicles/carriers were manually counted from maximum intensity z-projection of the entire z-stack confocal fluorescence microscopy image of the cell using ImageJ and used for further analysis. To quantify integrin signal at the basal membrane, a single plane with maximum basal signal was selected from a z-stack and used for downstream analyses. A total cell mask was obtained from a sum projection high contrast and used to analyze total intensity of the active integrin signal (Integrated Density). To measure FA area per cell, a total cell mask was obtained from a sum projection with high contrast. A binary image from the FA marker was used to define FA area in the total cell mask. To quantify pMLC signal at basal cell-substrate interface, a single plane with maximum basal signal was selected from a z-stack and used for downstream analyses. A total cell mask was obtained from phalloidin staining and used to analyze total intensity of pMLC signal (Integrated Density).

### Statistical analysis and data representation

Data were visualized and represented graphically using SuperPlots (Lord et al., 2020), as indicated in the corresponding figure panels. In brief, each small, light-colored symbol represents one individual cell measurement (N individual measurements per replicate), whereas the larger, black-outlined colored symbols represent the mean of all cells measured within one independent biological replicate (n replicates). Different colors denote distinct biological replicates. Horizontal black lines and error bars represent the mean and standard error of the mean (SEM), respectively, across biological replicates, thereby capturing variability between independent experiments. Individual cell measurements (N) are displayed to illustrate within-experiment variability, but were not treated as independent n (number of biological replicates) for statistical testing. Statistical analyses were performed in GraphPad Prism (version 11.0). P-values are reported as exact numerical values, and the statistical tests used are indicated in the figure legends. All quantification data underlying SuperPlot-based figures are available in Zenodo at doi.org/10.5281/zenodo.21073559. The repository includes GraphPad Prism project files used for plotting and statistical analysis.

### Online supplemental material

***Fig. S1*** shows experiments related to spatial exocytosis of CARTS close to FAs. ***Fig. S2*** is related to ***Fig. 1*** and shows additional results indicating that adhesion-dependent cell states associate with CARTS formation. ***Fig. S3*** shows experiments indicating that early cell spreading on FN transiently increases pMLC levels. ***Fig. S4*** is related to ***Fig. 5***, and shows additional experiments and information on MT hyperacetylation upon tubacin treatment. ***Fig. S5*** is related to ***Fig. 8***, and provides additional experiments on how Golgi-derived export is necessary for cell spreading and mechanoadaptation.

## DATA AVAILABILITY

The data underlying all figures are available in the published article and its online supplemental material. All quantification data underlying SuperPlot-based figures are available in Zenodo at doi.org/10.5281/zenodo.21073559. The repository includes GraphPad Prism project files used for plotting and statistical analysis.

## SUPPLEMENTARY FIGURE LEGENDS

**Figure S1. CARTS are delivered close to FAs.**

**(A)** TIRF image of a live HeLa cell expressing Paxillin-eGFP (green) and mKate2-FM4-PAUF (magenta). White squares highlight (i) FA-enriched (FA+) and (ii) FA-non-enriched (FA-) regions within the PM of the cell. Zoom-ins of the highlighted regions show a frame time sequence of the two merged channels. White circles highlight the vesicle disappearance along the frame sequence (i), and a random position of equal size in the FA- region of the cell. Plots on the right correspond to the fluorescence intensity profiles of mKate2-FM4-PAUF signal measured within the highlighted boxed regions show in the respective zoom-ins. Scale bars are 10 µm (main image), and 1 µm (zoom-ins). (**B**) Knockdown efficiency of ELKS in HeLa cells stably expressing PAUF-MycHis at 72 h after siRNA transfection. (**C**) Distribution of CARTS upon control and ELKS knockdown in HeLa cells stably expressing PAUF-MycHis. The cells plated on FN-coated coverslips were fixed and visualized with an anti-Myc monoclonal antibody. Scale bar, 10 μm. (**D**) Accumulation of CARTS near FAs. The cells plated on FN-coated coverslips were fixed and visualized with an anti-Myc polyclonal antibody and an anti-Paxillin monoclonal antibody (clone 349). High magnifications of the boxed areas are shown in the right of each panel. Scale bars, 10 μm. **(E)** HeLa cell expressing Paxillin-eGFP (green), and mKate2-FM4-PAUF (magenta), subjected to an SPI assay (see Methods), fixed at the indicated times after cargo release from the ER, and imaged by confocal fluorescence microscopy. Schematic of the experiment is shown on top. D/D is D/D solubilizer, CHX is cycloheximide. Scale bar, 10µm. (**F**) Quantification of (**E**), showing a 2D histogram (and corresponding 1D projections on each cartesian axis) of mKate2-FM4-PAUF fluorescence intensity per pixel as a function of the signed distance from that pixel to the closest FA area (nearest neighbor distance, see Methods for details), at the indicated time points. Source data are available for this figure: SourceData FS1.

**Figure S2. Adhesion-dependent cell states associate with CARTS formation.**

(**A**) Confocal fluorescence microscopy images of fixed HeLa cells seeded on FN or PLL for the indicated time points and immunostained against endogenous paxillin. Zoom-ins of the highlighted regions are shown on top. Scale bar, 10 µm. (**B**) SuperPlot showing the number of CARTS per cell (small, light-colored symbols; N∼10 per biological replicate) on cells lifted using Trypsin or EDTA (see Methods). The mean value for each independent biological replicate (larger, black-outlined circles; n=3). A repeated-measures 2-way ANOVA test was performed, and P values were obtained using Tukey’s post-hoc multiple comparison test, with only the values for the comparisons between Trypsin and EDTA being shown. (**C**) Spinning disk confocal fluorescence microscopy images of fixed HeLa cells that were plated on stiff (30 kPa) and soft (2 kPa) PAA gels, let spread for 4 h, fixed, and immunostained for vinculin, a FA marker. Scale bars, 10 µm. (**D**) Confocal microscopy images of fixed HeLa cells plated on FN-coated glass coverslips, or on stiff (30 kPa) or soft (2 kPa) FN-coated PAA gels. 4h after plating them, cells were fixed and processed for immunofluorescence microscopy against endogenous active ß1 integrin (9EG7 antibody) and paxillin. (**E, F**) SuperPlots showing quantification of the fluorescence intensity signal of active ß1 integrin per cell (**E**) and of the FA area per cell (**F**) from cells in (**D**). A repeated-measures 1-way ANOVA test was performed using Tukey’s post-hoc multiple comparison test (n=3 replicates, N∼10 cells per replicate).

**Figure S3: Early cell spreading on FN transiently increases pMLC levels.**

(**A**) Representative confocal microscopy images of HeLa cells seeded on FN or PLL for the indicated times, fixed, and stained for pMLC (pMLC Ser19-specific antibody) and F-actin (using Phalloidin-Alexa Fluor 647). Scale bars, 10 µm. **(B)** SuperPlot of the quantification of pMLC signal at the basal cell-substrate interface from cells treated as in (**A**). A repeated-measures 2-way ANOVA test was performed. P values using Tukey’s post-hoc multiple comparison test are reported (N∼10 cells per biological replicate; n=3 biological replicates).

**Figure S4. MT hyperacetylation upon tubacin treatment.**

(**A**) Representative confocal microscopy images of fixed HeLa cells stained for acetylated or total tubulin subjected or not to a treatment with 10 µM tubacin. Cell contours are indicated by white curves. Scale bar, 10 µm. (**B**) SuperPlots showing the ratio between fluorescence intensity of acetylated vs. total tubulin following the different treatments (n=3 replicates; N∼10 cells per replicate). A two-sided parametric ratio paired t-test was used. The P value is indicated in the plot. **(C)** Schematics of the pipeline followed in the tubacin-induced stimulation of the secretory pathway.

**Figure S5: Golgi-derived export is necessary for cell spreading and mechanoadaptation.**

(**A**) Representative confocal microscopy images of fixed HeLa cells transfected with the GFP-mem plasmid, subjected or not to a treatment with 10 µM Golgicide A. Scale bars, 10 µm; representative cell length scales shown. (**B**) SuperPlots showing individual cell measurements (small, light-colored symbols; N∼10 per biological replicate) and the mean value for each independent biological replicate (larger, black-outlined circles; n=3). Each color represents a different experimental replicate. A repeated-measures 2-way ANOVA test using Fisher’s LSD test was performed. (**C**) Confocal microscopy images of HeLa cells pre-seeded on FN-coated glass coverslips for 30 min and then treated with CRT0066101 (5 µM) or DMSO for 30 min (total time 1h) or 210 min (total time 4h), fixed and processed for immunofluorescence microscopy against endogenous active ß1 integrin (9EG7 antibody) and paxillin. Scale bar, 10 µm. (**D, E**) SuperPlots showing quantification of the fluorescence intensity signal of active ß1 integrin per cell (**D**) and of the FA area per cell (**E**) from cells in (**C**). A repeated-measures 2-way ANOVA test was performed using Tukey’s post-hoc multiple comparison test (N=3 replicates, n∼10 cells per replicate). (**F**) Confocal microscopy images of HeLa cells pre-seeded on FN-coated glass coverslips for 30 min and then treated with CRT0066101 (5 µM) or DMSO for 210 min (total time 4h), fixed and processed for immunofluorescence microscopy against acetylated and total α-tubulin. Scale bar, 10 µm. (**G**) SuperPlot showing quantification of the acetylated vs. total α-tubulin intensity signal per cell, from cells in (**F**). A repeated-measures 2-way ANOVA test was performed using Tukey’s post-hoc multiple comparison test (N=3 replicates, n∼10 cells per replicate). A two-sided parametric ratio paired t-test was used. The P value is indicated in the plot.

**Video 1. Synchronized release of mKate2-FM4-PAUF.** HeLa cells transiently expressing mKate2-FM4-PAUF and Paxillin-eGFP were observed live by TIRF microscopy. Paxillin images (not shown in the video, see ***Fig. S1A***) were taken as a reference of FAs before acquiring synchronized cargo export. Acquisition of PAUF channel (grayscale signal) was performed at 1 fps for 45 min, and sped up 50-fold. D/D solubilizer was added right before imaging started. Scale bar, 10 µm.

## ACKNOWLEDGMENTS

We thank members of the Single Molecule Biophotonics lab at ICFO, the Intracellular Transport: Engineering and Mechanisms Team at Institut Curie, the Intracellular Mechanics group and the Physics of Living Systems team at MSC, as well as Neus Sanfeliu-Cerdán, Santosh Phuyal, Hesso Farhan, Aurélien Roux, Gaëlle Boncompain, Pierre Sens, and Vivek Malhotra for valuable discussions. We thank Merche Rivas, Marina Perez, Angel Sandoval, and Maria Marsal for technical support at ICFO. We thank Angelika Hausser (University of Stuttgart) for kindly sharing the HeLa Flp-IN T-Rex GPKDrep cell line and the phospho-serine 294 (pS294)-specific antibody. We thank Rick Horwitz, Francesc Tebar, Vivek Malhotra, and Stefan Matile for kindly sharing reagents. We acknowledge support from the Government of Spain (RYC-2017-22227, PID2019-106232RB-I00/10.13039/501100011033/110198RB-I00, PID2020-113068RB-I00/10.13039/501100011033 and PID2023-147711NB-100; PID2022-138282NB-I00 project funded by the MCIN/AEI/10.13039/501100011033/FEDER, UE; PID2022-142672NB-I00; “Unidad de Excelencia María de Maeztu” CEX2024-001431-M, funded by MICIU/AEI/10.13039/501100011033 to MELIS-UPF, and Severo Ochoa CEX2019-000910-S to ICFO and CEX2023-001282-S to IBEC), Fundació Privada Cellex, Fundació Privada Mir-Puig, and Generalitat de Catalunya (CERCA, AGAUR), ERC Advanced Grants NANO-MEMEC (GA 788546) and MechanoSynth (GA 101097753), as well as LaserLab 4 Europe (GA 654148). E.A. acknowledges the support of the Beatriu de Pinós postdoctoral fellowship program of the Department of Research and Universities of the Generalitat de Catalunya (2023 BP 00210). N.M. acknowledges funding from the European Union H2020 under Marie Sklodowska-Curie grant 754558-PREBIST. J. A.-C. acknowledges funding from the European Union H2020 under the Marie Sklodowska-Curie grant agreement No 847517. A.W. was supported by joint funding from an ICFO Student Research Fellowship (Fall 2024) and a grant from Homerton College, Cambridge, UK. Y. W. acknowledges support from Grants-in-Aid for Scientific Research from the Ministry of Education, Culture, Sports, Science, and Technology of Japan (grant number 25K09568); and AMED Multidisciplinary Frontier Brain and Neuroscience Discoveries (Brain/MINDS 2.0) (grant number JP24wm0625506). P.R.-C. acknowledges support from the prize “ICREA Academia” for excellence in research. This work was supported by the French Agence Nationale de la Recherche (ANR), grant number ANR-22-CE13-0044 ‘MECHANGOLGI’ project. C.B.N. was supported by the international EuReCa PhD program of Institut Curie, H2020-MSCA-COFUND-2018-EuReCa-Grant Agreement number: 847718.

## REFERENCES

Aceiton, P. 2025. B cell mechanotransduction via ATAT1 coordinates actin and lysosomal dynamics at the immune synapse. Journal of Cell Biology. 224:e202407181. doi:10.1083/jcb.202407181.

Ahmed, W.W., and T.A. Saif. 2014. Active transport of vesicles in neurons is modulated by mechanical tension. Sci. Rep. 4:4481. doi:10.1038/srep04481.

Amar, K., F. Wei, J. Chen, and N. Wang. 2021. Effects of forces on chromatin. APL Bioeng. 5:41503. doi:10.1063/5.0065302.

Apodaca, G. 2002. Modulation of membrane traffic by mechanical stimuli. Am. J. Physiol. Renal Physiol. 282:F179–F190. doi:10.1152/ajprenal.2002.282.2.f179.

Barber-Perez, N., M. Georgiadou, C. Guzman, A. Isomursu, H. Hamidi, and J. Ivaska. 2020. Mechano-responsiveness of fibrillar adhesions on stiffness-gradient gels. J. Cell Sci. 133:jcs242909. doi:10.1242/jcs.242909.

Bazzoni, G., D.T. Shih, C.A. Buck, and M.E. Hemler. 1995. Monoclonal antibody 9EG7 defines a novel β1 integrin epitope induced by soluble ligand and manganese, but inhibited by calcium. Journal of Biological Chemistry. 270:25570–25577. doi:10.1074/JBC.270.43.25570/ASSET/66E9BD63-EC30-4569-99CD-AE653DC44039/MAIN.ASSETS/GR9.JPG.

Boncompain, G., S. Divoux, N. Gareil, H. De Forges, A. Lescure, L. Latreche, V. Mercanti, F. Jollivet, G. Raposo, and F. Perez. 2012. Synchronization of secretory protein traffic in populations of cells. Nat. Methods. 9:493–498. doi:10.1038/nmeth.1928.

Borges, S., E.A. Perez, E.A. Thompson, D.C. Radisky, X.J. Geiger, and P. Storz. 2015. Effective Targeting of Estrogen Receptor-Negative Breast Cancers with the Protein Kinase D Inhibitor CRT0066101. Mol. Cancer Ther. 14:1306–1316. doi:10.1158/1535-7163.mct-14-0945.

Boulant, S., C. Kural, J.-C. Zeeh, F. Ubelmann, and T. Kirchhausen. 2011. Actin dynamics counteract membrane tension during clathrin-mediated endocytosis. Nat. Cell Biol. 13:1124–1131. doi:10.1038/ncb2307.

Campelo, F., and V. Malhotra. 2012. Membrane fission: The biogenesis of transport carriers. Annu. Rev. Biochem. 81:407–427. doi:10.1146/annurev-biochem-051710-094912.

Carlsson, A.E. 2018. Membrane bending by actin polymerization. Curr. Opin. Cell Biol. 50:1–7. doi:10.1016/J.CEB.2017.11.007.

Chakraborty, A., S.M. Pitke, B.R. Rajeshwari, A. Dasgupta, N. Buwa, R. Behera, M. Jayakrishnan, and N. Balasubramanian. 2026. KIF5B and Dynein-Dependent Golgi Organization: Role in Adhesion-Dependent Microtubule Acetylation. Traffic. 27:e70033. doi:10.1111/TRA.70033.

Chastney, M.R., J. Kaivola, V.-M. Leppanen, and J. Ivaska. 2025. The role and regulation of integrins in cell migration and invasion. Nat. Rev. Mol. Cell Biol. 26:147–167. doi:10.1038/s41580-024-00777-1.

Chen, Z., P. Chen, J. Li, E. Landao-Bassonga, J. Papadimitriou, J. Gao, D. Liu, A. Tai, J. Ma, D. Lloyd, B.F. Kennedy, and M.H. Zheng. 2025. External strain on the plasma membrane is relayed to the endoplasmic reticulum by membrane contact sites and alters cellular energetics. Sci. Adv. 11:eads6132. doi:10.1126/sciadv.ads6132.

Cho, S., J. Irianto, and D.E. Discher. 2017. Mechanosensing by the nucleus: From pathways to scaling relationships. Journal of Cell Biology. 216:305–315. doi:10.1083/JCB.201610042.

Colom, A., E. Derivery, S. Soleimanpour, C. Tomba, M.D. Molin, N. Sakai, M. González-Gaitán, S. Matile, A. Roux, M. Gonzalez-Gaitan, S. Matile, and A. Roux. 2018. A fluorescent membrane tension probe. Nat. Chem. 10:1118–1125. doi:10.1038/s41557-018-0127-3.

Dai, J., and M.P. Sheetz. 1995. Regulation of Endocytosis, Exocytosis, and Shape by Membrane Tension. Cold Spring Harb. Symp. Quant. Biol. 60:567–571. doi:10.1101/SQB.1995.060.01.060.

Deb Roy, A., C.S. Gonzalez, M. Stanislauskas, F. Shahid, E. Yadav, J. Rezek, and T. Inoue. 2026. OptoTAT reveals microtubule acetylation as a rapid trigger for GEF-H1-mediated cell migration. J. Cell Biol. 225. doi:10.1083/JCB.202508095/281653.

Egea, G., F. Lazaro-Dieguez, and M. Vilella. 2006. Actin dynamics at the Golgi complex in mammalian cells. Curr. Opin. Cell Biol. 18:168–178. doi:10.1016/j.ceb.2006.02.007.

Eisler, S.A., F. Curado, G. Link, S. Schulz, M. Noack, M. Steinke, M.A. Olayioye, and A. Hausser. 2018. A Rho signaling network links microtubules to PKD controlled carrier transport to focal adhesions. Elife. 7:e35907. doi:10.7554/eLife.35907.

Farhan, H., I. Raote, F. Campelo, L. Ge, K. Hirschberg, A. Forrester, G. Zanetti, J. Lippincott-Schwartz, J.C. Pastor-Pareja, F. Perez, K. Saito, and V. Malhotra. 2025. Towards a unified framework for the function of endoplasmic reticulum exit sites. Nature Reviews Molecular Cell Biology 2025 26:12. 26:957–969. doi:10.1038/s41580-025-00899-0.

Fourriere, L., A.J. Jimenez, F. Perez, and G. Boncompain. 2020. The role of microtubules in secretory protein transport. J. Cell Sci. 133:jcs237016. doi:10.1242/jcs.237016.

Fourriere, L., A. Kasri, N. Gareil, S. Bardin, H. Bousquet, D. Pereira, F. Perez, B. Goud, G. Boncompain, and S. Miserey-Lenkei. 2019. RAB6 and microtubules restrict protein secretion to focal adhesions. Journal of Cell Biology. 218:2215–2231. doi:10.1083/jcb.201805002.

Fu, Y., and C.S. Rubin. 2011. Protein kinase D: Coupling extracellular stimuli to the regulation of cell physiology. EMBO Rep. 12:785–796. doi:10.1038/embor.2011.139.

Fuchs, Y.F., S.A. Eisler, G. Link, O. Schlicker, G. Bunt, K. Pfizenmaier, and A. Hausser. 2009. A Golgi PKD activity reporter reveals a crucial role of PKD in nocodazole-induced Golgi dispersal. Traffic. 10:858–867. doi:10.1111/j.1600-0854.2009.00918.x.

Fugmann, T., A. Hausser, P. Schöffler, S. Schmid, K. Pfizenmaier, and M.A. Olayioye. 2007. Regulation of secretory transport by protein kinase D-mediated phosphorylation of the ceramide transfer protein. Journal of Cell Biology. 178:15–22. doi:10.1083/jcb.200612017.

Gauthier, N.C., M.A. Fardin, P. Roca-Cusachs, and M.P. Sheetz. 2011. Temporary increase in plasma membrane tension coordinates the activation of exocytosis and contraction during cell spreading. Proceedings of the National Academy of Sciences. 108:14467–14472. doi:10.1073/pnas.1105845108.

Geiger, B., J.P. Spatz, and A.D. Bershadsky. 2009. Environmental sensing through focal adhesions. Nat. Rev. Mol. Cell Biol. 10:21–33. doi:10.1038/nrm2593.

Grigoriev, I., D. Splinter, N. Keijzer, P.S. Wulf, J. Demmers, T. Ohtsuka, M. Modesti, I. V. Maly, F. Grosveld, C.C. Hoogenraad, and A. Akhmanova. 2007. Rab6 Regulates Transport and Targeting of Exocytotic Carriers. Dev. Cell. 13:305–314. doi:10.1016/j.devcel.2007.06.010.

Grigoriev, I., K. Lou Yu, E. Martinez-Sanchez, A. Serra-Marques, I. Smal, E. Meijering, J. Demmers, J. Peränen, R.J. Pasterkamp, P. Van Der Sluijs, C.C. Hoogenraad, and A. Akhmanova. 2011. Rab6, Rab8, and MICAL3 cooperate in controlling docking and fusion of exocytotic carriers. Current Biology. 21:967–974. doi:10.1016/j.cub.2011.04.030.

Guet, D., K. Mandal, M. Pinot, J. Hoffmann, Y. Abidine, W. Sigaut, S. Bardin, K. Schauer, B. Goud, and J.-B. Manneville. 2014. Mechanical Role of Actin Dynamics in the Rheology of the Golgi Complex and in Golgi-Associated Trafficking Events. Current Biology. 24:1700–1711. doi:10.1016/j.cub.2014.06.048.

Guo, M., A.F. Pegoraro, A. Mao, E.H. Zhou, P.R. Arany, Y. Han, D.T. Burnette, M.H. Jensen, K.E. Kasza, J.R. Moore, F.C. Mackintosh, J.J. Fredberg, D.J. Mooney, J. Lippincott-Schwartz, and D.A. Weitz. 2017. Cell volume change through water efflux impacts cell stiffness and stem cell fate. Proceedings of the National Academy of Sciences. 114:E8618–E8627. doi:10.1073/pnas.1705179114.

Haggarty, S.J., K.M. Koeller, J.C. Wong, R.A. Butcher, and S.L. Schreiber. 2003a. Multidimensional Chemical Genetic Analysis of Diversity-Oriented Synthesis-Derived Deacetylase Inhibitors Using Cell-Based Assays. Chem. Biol. 10:383–396. doi:10.1016/s1074-5521(03)00095-4.

Haggarty, S.J., K.M. Koeller, J.C. Wong, C.M. Grozinger, and S.L. Schreiber. 2003b. Domain-selective small-molecule inhibitor of histone deacetylase 6 (HDAC6)-mediated tubulin deacetylation. Proceedings of the National Academy of Sciences. 100:4389–4394. doi:10.1073/pnas.0430973100.

Harikumar, K.B., A.B. Kunnumakkara, N. Ochi, Z. Tong, A. Deorukhkar, B. Sung, L. Kelland, S. Jamieson, R. Sutherland, T. Raynham, M. Charles, A. Bagherzadeh, C. Foxton, A. Boakes, M. Farooq, D. Maru, P. Diagaradjane, Y. Matsuo, J. Sinnett-Smith, J. Gelovani, S. Krishnan, B.B. Aggarwal, E. Rozengurt, C.R. Ireson, and S. Guha. 2010. A Novel Small-Molecule Inhibitor of Protein Kinase D Blocks Pancreatic Cancer Growth *In vitro* and *In vivo*. Mol. Cancer Ther. 9:1136–1146. doi:10.1158/1535-7163.MCT-09-1145.

Hausser, A., P. Storz, S. Märtens, G. Link, A. Toker, and K. Pfizenmaier. 2005. Protein kinase D regulates vesicular transport by phosphorylating and activating phosphatidylinositol-4 kinase IIIβ at the Golgi complex. Nat. Cell Biol. 7:880–886. doi:10.1038/ncb1289.

Huet-Calderwood, C., F. Rivera-Molina, D. V. Iwamoto, E.B. Kromann, D. Toomre, and D.A. Calderwood. 2017. Novel ecto-tagged integrins reveal their trafficking in live cells. Nat. Commun. 8:570. doi:10.1038/s41467-017-00646-w.

Huet-Calderwood, C., F.E. Rivera-Molina, D.K. Toomre, and D.A. Calderwood. 2023. Fibroblasts secrete fibronectin under lamellipodia in a microtubule- and myosin II–dependent fashion. Journal of Cell Biology. 222:e202204100. doi:10.1083/jcb.202204100.

Jahed, Z., and M.R. Mofrad. 2019. The nucleus feels the force, LINCed in or not! Curr. Opin. Cell Biol. 58:114–119. doi:10.1016/j.ceb.2019.02.012.

Jimenez-Rojo, N. 2025. Optical Control of Membrane Viscosity Modulates ER-to-Golgi Trafficking. ACS Cent. Sci. doi:10.1021/acscentsci.5c00606.

Joo, E.E., and K.M. Yamada. 2014. MYPT1 regulates contractility and microtubule acetylation to modulate integrin adhesions and matrix assembly. Nat. Commun. 5. doi:10.1038/NCOMMS4510.

Joshi, P., A. Saha, R. Malaviya, D. Panda, G. Mehta, M. Pattanayak, V. Singh, and N. Balasubramanian. 2026. AXL receptor tyrosine kinase regulates Golgi organization and function via an adhesion-Arf1 signalling axis in breast and lung cancer cell lines. Biol. Open. 15. doi:10.1242/BIO.062581.

Jung, J., M.M. Khan, J. Landry, A. Halavatyi, P. Machado, M. Reiss, and R. Pepperkok. 2022. Regulation of the COPII secretory machinery via focal adhesions and extracellular matrix signaling. Journal of Cell Biology. 221:e202110081. doi:10.1083/jcb.202110081.

Kanchanawong, P., and D.A. Calderwood. 2023. Organization, dynamics and mechanoregulation of integrin-mediated cell–ECM adhesions. Nat. Rev. Mol. Cell Biol. 24:142–161. doi:10.1038/s41580-022-00531-5.

Kaverina, I., K. Rottner, and J.V. Small. 1998. Targeting, Capture, and Stabilization of Microtubules at Early Focal Adhesions. Journal of Cell Biology. 142:181–190. doi:10.1083/JCB.142.1.181.

Kechagia, J.Z., J. Ivaska, and P. Roca-Cusachs. 2019. Integrins as biomechanical sensors of the microenvironment. Nat. Rev. Mol. Cell Biol. 20:457–473. doi:10.1038/s41580-019-0134-2.

Kim, S.A., Y. Lee, D.E. Jung, K.H. Park, J.Y. Park, J. Gang, S.B. Jeon, E.C. Park, Y.G. Kim, B. Lee, Q. Liu, W. Zeng, S. Yeramilli, S. Lee, S.S. Koh, and S.Y. Song. 2009. Pancreatic adenocarcinoma up-regulated factor (PAUF), a novel up-regulated secretory protein in pancreatic ductal adenocarcinoma. Cancer Sci. 100:828–836. doi:10.1111/j.1349-7006.2009.01106.x.

Kim, T.J., C. Joo, J. Seong, R. Vafabakhsh, E.L. Botvinick, M.W. Berns, A.E. Palmer, N. Wang, T. Ha, E. Jakobsson, J. Sun, and Y. Wang. 2015. Distinct mechanisms regulating mechanical force-induced Ca2+ signals at the plasma membrane and the ER in human MSCs. Elife. 2015. doi:10.7554/ELIFE.04876.

Kirby, T.J., and J. Lammerding. 2018. Emerging views of the nucleus as a cellular mechanosensor. Nat. Cell Biol. 20:373–381. doi:10.1038/S41556-018-0038-Y.

Kosmalska, A.J., L. Casares, A. Elosegui-Artola, J.J. Thottacherry, R. Moreno-Vicente, V. González-Tarragó, M.Á. Del Pozo, S. Mayor, M. Arroyo, D. Navajas, X. Trepat, N.C. Gauthier, and P. Roca-Cusachs. 2015. Physical principles of membrane remodelling during cell mechanoadaptation. Nat. Commun. 6:7292. doi:10.1038/ncomms8292.

Krendel, M., F.T. Zenke, and G.M. Bokoch. 2002. Nucleotide exchange factor GEF-H1 mediates cross-talk between microtubules and the actin cytoskeleton. Nat. Cell Biol. 4:294–301. doi:10.1038/ncb773.

Krylyshkina, O., K.I. Anderson, I. Kaverina, I. Upmann, D.J. Manstein, J. V Small, and D.K. Toomre. 2003. Nanometer targeting of microtubules to focal adhesions. Journal of Cell Biology. 161:853–859. doi:10.1083/jcb.200301102.

Lachuer, H., L. Le, S. Leveque-Fort, B. Goud, and K. Schauer. 2023. Spatial organization of lysosomal exocytosis relies on membrane tension gradients. Proceedings of the National Academy of Sciences. 120:e2207425120. doi:10.1073/pnas.2207425120.

Laukaitis, C.M., D.J. Webb, K. Donais, and A.F. Horwitz. 2001. Differential Dynamics of alpha5 Integrin, Paxillin, and alpha-Actinin during Formation and Disassembly of Adhesions in Migrating Cells. Journal of Cell Biology. 153:1427–1440. doi:10.1083/jcb.153.7.1427.

Lerche, M., M. Mathieu, H. Hamidi, M. Chastney, G. Jacquemet, B.M.H. Bruininks, S. Kaptan, L. Malerød, N.M. Pedersen, A. Brech, N. Matoba, Y. Sato, I. Vattulainen, P. Roca-Cusachs, F. Perez, G. Boncompain, S. Miserey, and J. Ivaska. 2026. Regulation of cell dynamics by rapid integrin transport through the biosynthetic pathway. J. Cell Biol. 225. doi:10.1083/JCB.202508155/278539.

Li, Y., O. Kučera, D. Cuvelier, D.M. Rutkowski, M. Deygas, D. Rai, T. Pavlovič, F.N. Vicente, M. Piel, G. Giannone, D. Vavylonis, A. Akhmanova, L. Blanchoin, and M. Théry. 2023. Compressive forces stabilize microtubules in living cells. Nature Materials 2023 22:7. 22:913–924. doi:10.1038/s41563-023-01578-1.

Lord, S.J., K.B. Velle, R.D. Mullins, and L.K. Fritz-Laylin. 2020. SuperPlots: Communicating reproducibility and variability in cell biology. Journal of Cell Biology. 219:e202001064. doi:10.1083/jcb.202001064.

Lujan, P., C. Garcia-Cabau, Y. Wakana, J. Vera Lillo, C. Rodilla-Ramírez, H. Sugiura, V. Malhotra, X. Salvatella, M.F. Garcia-Parajo, and F. Campelo. 2024. Sorting of secretory proteins at the trans-Golgi network by human TGN46. Elife. 12. doi:10.7554/eLife.91708.

Maeda, Y., G. V. Beznoussenko, J. Van Lint, A.A. Mironov, V. Malhotra, J. Van Lint, A.A. Mironov, and V. Malhotra. 2001. Recruitment of protein kinase D to the trans-Golgi network via the first cysteine-rich domain. EMBO J. 20:5982–5990. doi:10.1093/emboj/20.21.5982.

Merta, H., K. Gov, T. Isogai, B. Paul, A. Sannigrahi, A. Radhakrishnan, G. Danuser, and W.M. Henne. 2025. Spatial proteomics of ER tubules reveals CLMN, an ER-actin tether at focal adhesions that promotes cell migration. Cell Rep. 44:115502. doi:10.1016/j.celrep.2025.115502.

Miserey-Lenkei, S., H. Bousquet, O. Pylypenko, S. Bardin, A. Dimitrov, G. Bressanelli, R. Bonifay, V. Fraisier, C. Guillou, C. Bougeret, A. Houdusse, A. Echard, and B. Goud. 2017. Coupling fission and exit of RAB6 vesicles at Golgi hotspots through kinesin-myosin interactions. Nat. Commun. 8:1–13. doi:10.1038/s41467-017-01266-0.

Miserey-Lenkei, S., G. Chalancon, S. Bardin, E. Formstecher, B. Goud, and A. Echard. 2010. Rab and actomyosin-dependent fission of transport vesicles at the golgi complex. Nat. Cell Biol. 12:645–654. doi:10.1038/ncb2067.

Moreno-Layseca, P., J. Icha, H. Hamidi, and J. Ivaska. 2019. Integrin trafficking in cells and tissues. Nat. Cell Biol. 21:122–132. doi:10.1038/s41556-018-0223-z.

Naughton, B.S., S.C. Devarkar, V. Todorow, S. Mallik, S. Oxendine, S. Junnarkar, Y. Ren, J. Berro, J. Kirstein, Y. Xiong, and C. Schlieker. 2025. Nodal modulator (NOMO) is a force-bearing transmembrane protein required for muscle differentiation. Journal of Cell Biology. 224:e202505010. doi:10.1083/jcb.202505010.

Pelham, R.J., and Y. Wang. 1997. Cell locomotion and focal adhesions are regulated by substrate flexibility. Proceedings of the National Academy of Sciences. 94:13661–13665. doi:10.1073/pnas.94.25.13661.

Pérez-González, C., G. Ceada, F. Greco, M. Matejčić, M. Gómez-González, N. Castro, A. Menendez, S. Kale, D. Krndija, A.G. Clark, V.R. Gannavarapu, A. Álvarez-Varela, P. Roca-Cusachs, E. Batlle, D.M. Vignjevic, M. Arroyo, and X. Trepat. 2021. Mechanical compartmentalization of the intestinal organoid enables crypt folding and collective cell migration. Nature Cell Biology 2021 23:7. 23:745–757. doi:10.1038/s41556-021-00699-6.

Phuyal, S., and F. Baschieri. 2020. Endomembranes: Unsung Heroes of Mechanobiology? Front. Bioeng. Biotechnol. 8:597721. doi:10.3389/fbioe.2020.597721.

Phuyal, S., E. Djaerff, A. Le Roux, M.J. Baker, D. Fankhauser, S.J. Mahdizadeh, V. Reiterer, A. Parizadeh, E. Felder, J.C. Kahlhofer, D. Teis, M.G. Kazanietz, S. Geley, L. Eriksson, P. Roca Cusachs, and H. Farhan. 2022. Mechanical strain stimulates COPII dependent secretory trafficking via Rac1. EMBO J. 41:e110596. doi:10.15252/embj.2022110596.

Pontes, B., P. Monzo, L. Gole, A.-L.L. Roux, A.J. Kosmalska, Z.Y. Tam, W. Luo, S. Kan, V. Viasnoff, P. Roca-Cusachs, L. Tucker-Kellogg, and N.C. Gauthier. 2017. Membrane tension controls adhesion positioning at the leading edge of cells. Journal of Cell Biology. 216:2959–2977. doi:10.1083/jcb.201611117.

Rawal, S., P. Keshavanarayana, D. Manoj, P. Khuntia, S. Banerjee, B. Thurakkal, R. Marwaha, F. Spill, and T. Das. 2025. Edge curvature drives endoplasmic reticulum reorganization and dictates epithelial migration mode. Nat. Cell Biol. 1–16. doi:10.1038/s41556-025-01729-3.

Ren, W.-W., R. Kawahara, K.G.N. Suzuki, P. Dipta, G. Yang, M. Thaysen-Andersen, and M. Fujita. 2025. MYO18B promotes lysosomal exocytosis by facilitating focal adhesion maturation. Journal of Cell Biology. 224:e202407068. doi:10.1083/jcb.202407068.

Ren, X.D., R. Wang, Q. Li, L.A.F. Kahek, K. Kaibuchi, and R.A.F. Clark. 2004. Disruption of Rho signal transduction upon cell detachment. J. Cell Sci. 117:3511–3518. doi:10.1242/JCS.01205.

Robinson, I.M., J.M. Finnegan, J.R. Monck, R.M. Wightman, and J.M. Fernandez. 1995. Colocalization of calcium entry and exocytotic release sites in adrenal chromaffin cells. Proceedings of the National Academy of Sciences. 92:2474–2478. doi:10.1073/PNAS.92.7.2474.

Romani, P., I. Brian, G. Santinon, A. Pocaterra, M. Audano, S. Pedretti, S. Mathieu, M. Forcato, S. Bicciato, J.B. Manneville, N. Mitro, and S. Dupont. 2019. Extracellular matrix mechanical cues regulate lipid metabolism through Lipin-1 and SREBP. Nat. Cell Biol. 21:338–347. doi:10.1038/s41556-018-0270-5.

Le Roux, A.-L., C. Tozzi, N. Walani, X. Quiroga, D. Zalvidea, X. Trepat, M. Staykova, M. Arroyo, and P. Roca-Cusachs. 2021. Dynamic mechanochemical feedback between curved membranes and BAR protein self-organization. Nat. Commun. 12:6550. doi:10.1038/s41467-021-26591-3.

Le Roux, A.L., V. Venturini, M. Gómez-González, A.E.M. Beedle, X. Quiroga, X. Menino, X. Trepat, and P. Roca-Cusachs. 2025. Equibiaxial Stretching Device for High Magnification Live-Cell Confocal Fluorescence Microscopy. Journal of Visualized Experiments. 2025-June. doi:10.3791/67520.

Saenz, J.B., W.J. Sun, J.W. Chang, J. Li, B. Bursulaya, N.S. Gray, and D.B. Haslam. 2009. Golgicide A reveals essential roles for GBF1 in Golgi assembly and function. Nat. Chem. Biol. 5:157–165. doi:10.1038/nchembio.144.

Saha, A., T. Sherkhane, and N. Balasubramanian. 2026. Differential AXL expression and Arf1 regulation control stiffness-dependent Golgi organization in breast cancer cells. J. Cell Sci. 139. doi:10.1242/JCS.263956.

Schmidt, C.J., and S.J. Stehbens. 2024. Microtubule control of migration: Coordination in confinement. Curr. Opin. Cell Biol. 86:102289. doi:10.1016/j.ceb.2023.102289.

Seetharaman, S., and S. Etienne-Manneville. 2019. Microtubules at focal adhesions - a double-edged sword. J. Cell Sci. 132:jcs232843. doi:10.1242/jcs.232843.

Seetharaman, S., B. Vianay, V. Roca, A.J. Farrugia, C. De Pascalis, B. Boeda, F. Dingli, D. Loew, S. Vassilopoulos, A. Bershadsky, M. Thery, and S. Etienne-Manneville. 2022. Microtubules tune mechanosensitive cell responses. Nat. Mater. 21:366–377. doi:10.1038/s41563-021-01108-x.

Serafino, G., S. Forciniti, E. Scarpa, A. Ranieri, L. Santorelli, G. De Blasi, S. Costantini, E.J. Lee, A. Galeone, A. Calcagnì, M. Pirozzi, L.L. del Mercato, R. Venditti, G. D’Angelo, S. Parashuraman, T. Verri, G. Gigli, E.S. Sztul, P. Grumati, L. Rizzello, D. Russo, and R. Rizzo. 2026. Mechanical Cues Regulate Cargo Sorting and Export at the Golgi. Advanced Science. 13:e06241. doi:10.1002/advs.202506241.

Singh, V., C. Erady, and N. Balasubramanian. 2018. Cell-matrix adhesion controls Golgi organization and function through Arf1 activation in anchorage-dependent cells. J. Cell Sci. 131:jcs215855. doi:10.1242/jcs.215855.

Song, Y., Z. Zhao, L. Xu, P. Huang, J. Gao, J. Li, X. Wang, Y. Zhou, J. Wang, W. Zhao, L. Wang, C. Zheng, B. Gao, L. Jiang, K. Liu, Y. Guo, X. Yao, and L. Duan. 2024. Using an ER-specific optogenetic mechanostimulator to understand the mechanosensitivity of the endoplasmic reticulum. Dev. Cell. 59:1396–1409.e5. doi:10.1016/j.devcel.2024.03.014.

Stehbens, S.J., M. Paszek, H. Pemble, A. Ettinger, S. Gierke, and T. Wittmann. 2014. CLASPs link focal-adhesion-associated microtubule capture to localized exocytosis and adhesion site turnover. Nat. Cell Biol. 16:558–570. doi:10.1038/ncb2975.

Strakova, K., A.I. Poblador-Bahamonde, N. Sakai, and S. Matile. 2019. Fluorescent Flipper Probes: Comprehensive Twist Coverage. Chemistry - A European Journal. 25:14935–14942. doi:10.1002/chem.201903604.

Sugiura, H., M. Fujii, Y. Terashima, Y. Takagi, J. Angulo-Capel, M. Tagaya, H. Inoue, K. Arasaki, F. Campelo, and Y. Wakana. 2026. A PKD-caveolin axis drives secretory carrier biogenesis at the TGN. bioRxiv. 2026.01.13.699385. doi:10.64898/2026.01.13.699385.

Théry, M. 2010. Micropatterning as a tool to decipher cell morphogenesis and functions. J. Cell Sci. 123:4201–4213. doi:10.1242/JCS.075150.

Thorpe, S.D., and D.A. Lee. 2017. Dynamic regulation of nuclear architecture and mechanics-a rheostatic role for the nucleus in tailoring cellular mechanosensitivity. Nucleus. 8:287–300. doi:10.1080/19491034.2017.1285988.

Torra, J., F. Campelo, and M.F. Garcia-Parajo. 2024. Tensing Flipper: Photosensitized Manipulation of Membrane Tension, Lipid Phase Separation, and Raft Protein Sorting in Biological Membranes. J. Am. Chem. Soc. 146:24114–24124. doi:10.1021/jacs.4c08580.

Totsukawa, G., Y. Yamakita, S. Yamashiro, D.J. Hartshorne, Y. Sasaki, and F. Matsumura. 2000. Distinct Roles of Rock (Rho-Kinase) and Mlck in Spatial Regulation of Mlc Phosphorylation for Assembly of Stress Fibers and Focal Adhesions in 3t3 Fibroblasts. Journal of Cell Biology. 150:797–806. doi:10.1083/JCB.150.4.797.

Venkova, L., A.S. Vishen, S. Lembo, N. Srivastava, B. Duchamp, A. Ruppel, A. Williart, S. Vassilopoulos, A. Deslys, J.M. Garcia Arcos, A. Diz-Muñoz, M. Balland, J.F. Joanny, D. Cuvelier, P. Sens, and M. Piel. 2022. A mechano-osmotic feedback couples cell volume to the rate of cell deformation. Elife. 11. doi:10.7554/ELIFE.72381.

Vidal-Quadras, M., M. Gelabert-Baldrich, D. Soriano-Castell, A. Llado, C. Rentero, M. Calvo, A. Pol, C. Enrich, and F. Tebar. 2011. Rac1 and Calmodulin Interactions Modulate Dynamics of ARF6-Dependent Endocytosis. Traffic. 12:1879–1896. doi:10.1111/j.1600-0854.2011.01274.x.

Wakana, Y., and F. Campelo. 2021. The PKD-Dependent Biogenesis of TGN-to-Plasma Membrane Transport Carriers. Cells. 10:1618. doi:10.3390/cells10071618.

Wakana, Y., J. Van Galen, F. Meissner, M. Scarpa, R.S. Polishchuk, M. Mann, and V. Malhotra. 2012. A new class of carriers that transport selective cargo from the trans Golgi network to the cell surface. EMBO J. 31:3976–3990. doi:10.1038/emboj.2012.235.

Wakana, Y., K. Hayashi, T. Nemoto, C. Watanabe, M. Taoka, J. Angulo-Capel, M.F. Garcia-Parajo, H. Kumata, T. Umemura, H. Inoue, K. Arasaki, F. Campelo, and M. Tagaya. 2021. The ER cholesterol sensor SCAP promotes CARTS biogenesis at ER–golgi membrane contact sites. Journal of Cell Biology. 220. doi:10.1083/jcb.202002150.

Wu, X.S., S. Elias, H. Liu, J. Heureaux, P.J. Wen, A.P. Liu, M.M. Kozlov, and L.G. Wu. 2017. Membrane Tension Inhibits Rapid and Slow Endocytosis in Secretory Cells. Biophys. J. 113:2406–2414. doi:10.1016/j.bpj.2017.09.035.

Yeaman, C., M.I. Ayala, J.R. Wright, F. Bard, C. Bossard, A. Ang, Y. Maeda, T. Seufferlein, I. Mellman, W.J. Nelson, and V. Malhotra. 2004. Protein kinase D regulates basolateral membrane protein exit from trans-Golgi network. Nat. Cell Biol. 6:106–112. doi:10.1038/ncb1090.

Yuan, T., J. Lu, J. Zhang, Y. Zhang, and L. Chen. 2015. Spatiotemporal Detection and Analysis of Exocytosis Reveal Fusion “Hotspots” Organized by the Cytoskeleton in Endocrine Cells. Biophys. J. 108:251–260. doi:10.1016/J.BPJ.2014.11.3462.

Zilberman, Y., N.O. Alieva, S. Miserey-Lenkei, A. Lichtenstein, Z. Kam, H. Sabanay, and A. Bershadsky. 2011. Involvement of the Rho-mDia1 pathway in the regulation of Golgi complex architecture and dynamics. Mol. Biol. Cell. 22:2900–2911. doi:10.1091/mbc.e11-01-0007.

