## Supplementary figures and images for "Mechanical forces stimulate Golgi export"

### Source Data Supp. Fig. S1

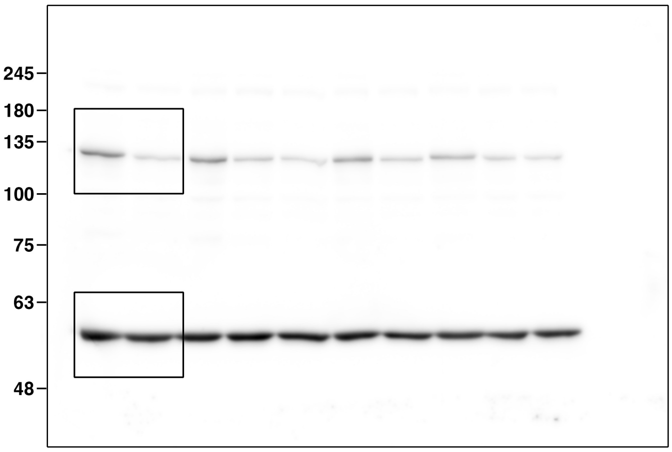

### Supplementary Figures

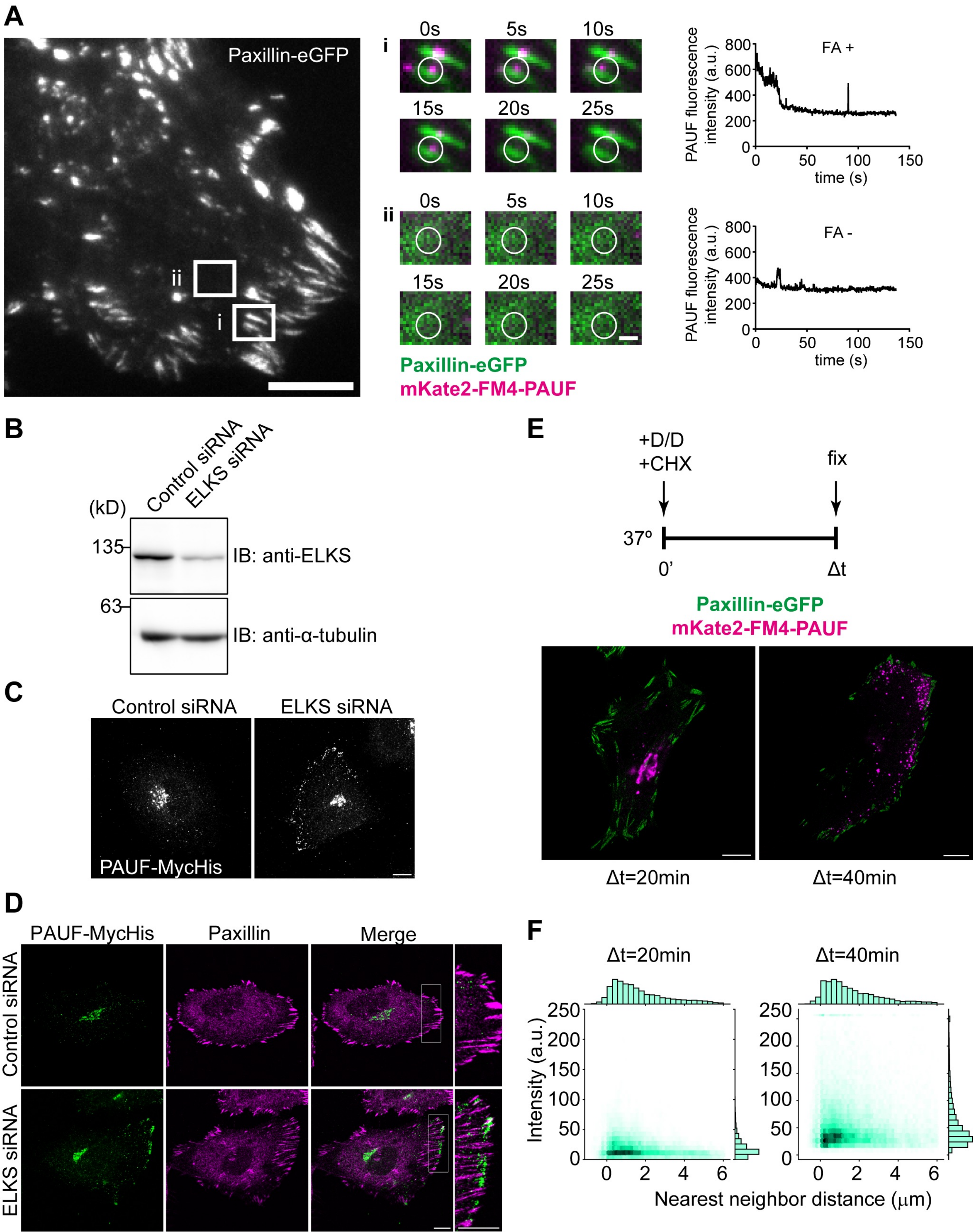

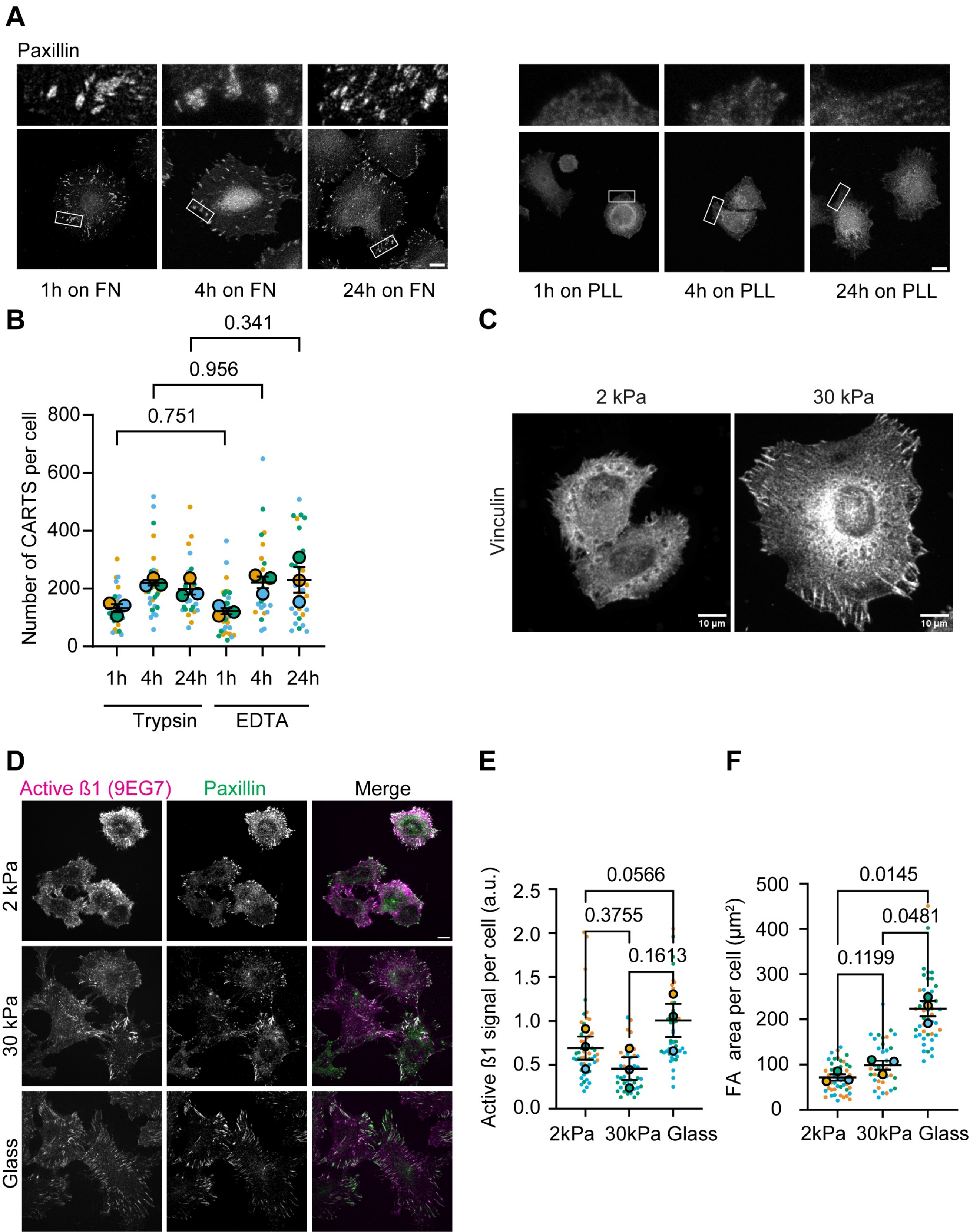

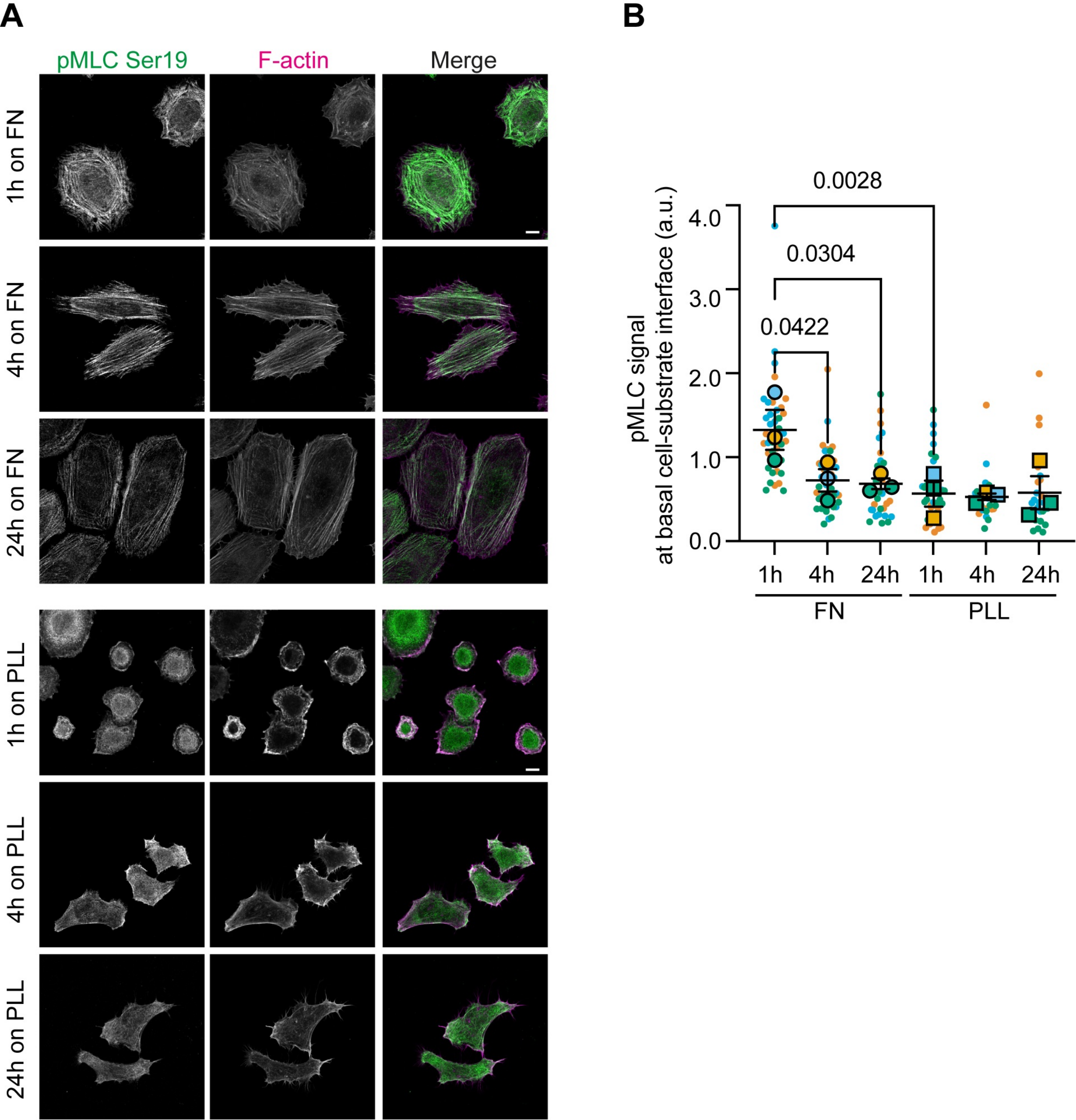

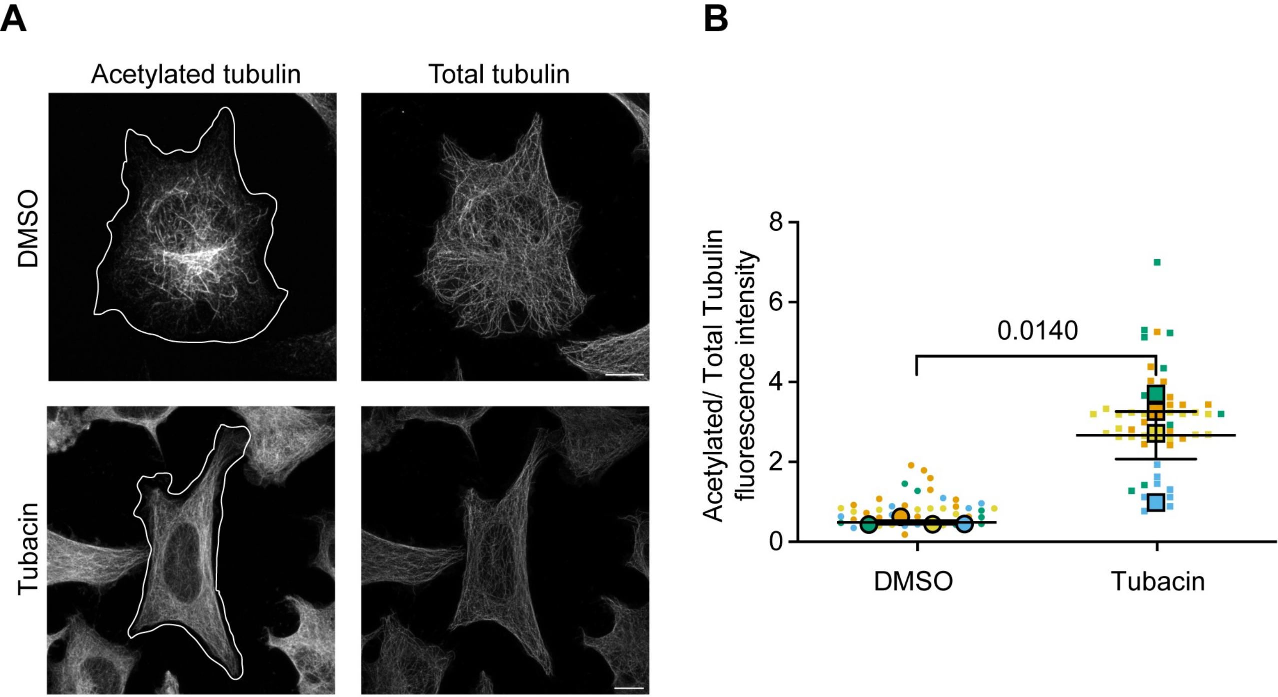

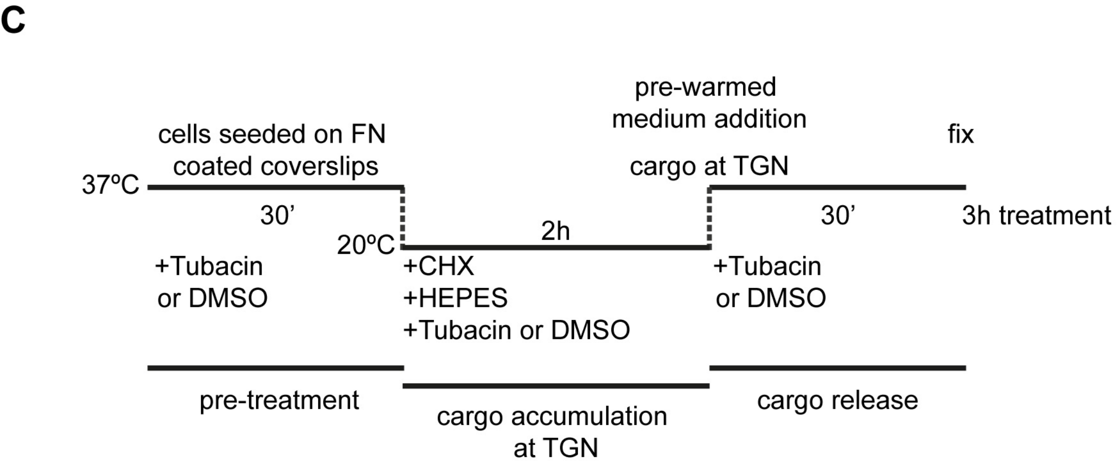

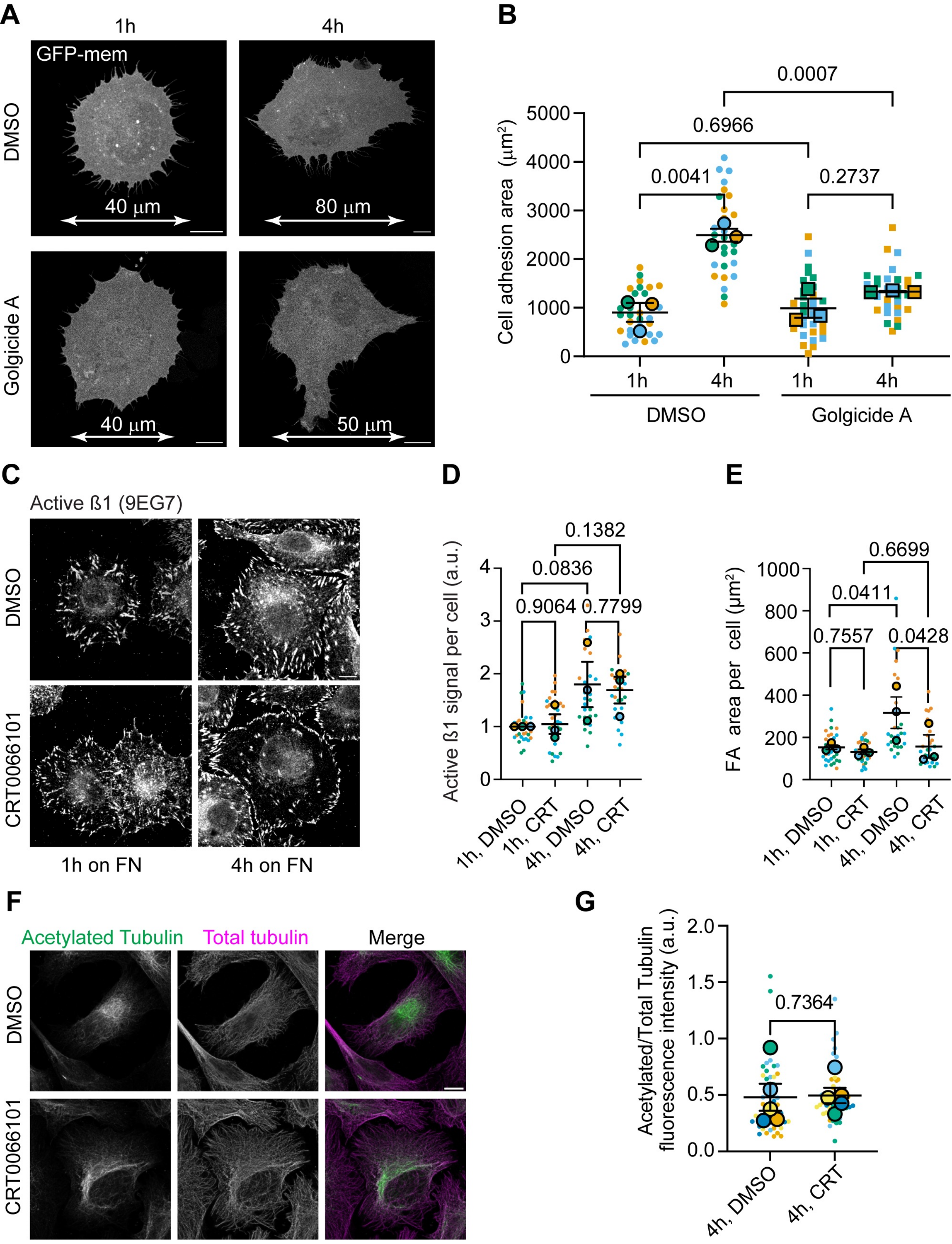
